# Metabolic Vulnerabilities of Temozolomide-Resistant Glioblastoma Cells: Implications for Targeted Therapies and Overcoming Chemoresistance

**DOI:** 10.1101/2024.03.01.582997

**Authors:** Meitham Amereh, Amir Seyfoori, Shahla Shojaei, Sarah Lane, Tian Zhao, Mahdieh Shokrollahi, Julian Lum, Patrick Walter, Mohsen Akbari

**Affiliations:** Laboratory for Innovations in Micro Engineering (LiME), Department of Mechanical Engineering, University of Victoria, Victoria, BC, V8P 5C2, Canada; Department of Human Anatomy and Cell Science, Max Rady College of Medicine, Rady Faculty of Health Sciences, University of Manitoba, Winnipeg, MB R3T 2N2, Canada; Department of Biology, Centre for Biomedical Research, University of Victoria, BC, Canada; Trev and Joyce Deeley Research Centre, BC Cancer, Victoria, BC V8R 6V5, Canada; Department of Biochemistry and Microbiology, University of Victoria, Victoria, BC V8W 2Y2, Canada; Hematology/Oncology, UCSF Benioff Children’s Hospital, Oakland, USA; Terasaki Institute for Biomedical Innovations, Los Angeles, CA 91367, USA

## Abstract

Chemoresistance is a major clinical challenge in the management of glioblastoma (GB), making it difficult to achieve long-term success with traditional treatments. Therefore, there is a need for the development of novel drugs. We explored the metabolic vulnerabilities of temozolomide (TMZ)-resistant GB and their potential implications for targeted therapies. In monolayer and tumoroid cultures, we found elevated reliance on oxidative phosphorylation in TMZ-resistant cells. Notably, iron reduction in TMZ-resistant cells reduced viability and proliferation, upregulated hypoxia-inducible factor 1-α (Hif1-α) expression, induced autophagy, inhibited autophagic flux, and increased reactive oxygen species (ROS) generation, indicating the significance of iron in metabolic vulnerabilities of these cells. Hypoxic cells showed acquired resistance to iron chelation compared to their normoxic state, suggesting an adaptive mechanism associated to hypoxia. Viability, size, and invasion were reduced in TMZ-resistant tumoroids. Additionally, we reported IC50 for the combination of TMZ with a range of DFO and DFP, making the combination therapy a promising drug candidate to improve therapeutic treatments.

**Teaser:** Combining iron reduction and chemotherapy in drug-resistant glioblastoma cells enhances therapeutic outcomes.

## Introduction

Glioblastoma (GB) is among the most malignant glial tumors with a low survival rate of 5.5% of patients living five years post diagnosis [1], [2]. Temozolomide (TMZ), the only FDA-approved drug for GB, is an alkylating agent that binds an alkyl group to DNA and causes insertion of thymine instead of a cytosine opposite the methylguanine during subsequent DNA replication, and hence leads to apoptosis [3]. The limited effectiveness of TMZ in glioblastoma treatment is attributed to the complex cellular responses, which include factors such as O^6^-methylguanine-DNA methyltransferase (MGMT), mutated P53, upregulated MCL-1 through transcriptional coactivator with PDZ-binding motif (TAZ), and the presence of tumor residues containing resistant cancer stem cell (CSC) populations [4]– [6]. In particular, chemoresistant cells exhibit distinct alterations in cell metabolism when compared to their non-resistant counterparts, including significant variations in glucose uptake, iron utilization, and responses to hypoxia within the tumor environment [7]. These deviations in metabolism, which enhance energy production, aid in the preservation of drug resistance [8].

Iron metabolism plays a significant role in promoting chemoresistance, as resistant cells can dysregulate iron homeostasis to foster tumor growth and evade treatment. Iron is one of the important contributors in redox reactions, such as adenosine triphosphate (ATP) production and respiration, as it is found in mitochondrial complex I and II [9]. It is known as an important part of prosthetic groups of proteins such as heme, iron-sulfur, cytochrome P450 and cytochromes, which are required for oxidative phosphorylation and detoxification processes [9]. ROS generation is also another important reactions (Fenton reaction) where iron’s capability of losing and gaining electrons marks it as one of the key players inducing cell toxicity, knows as ferroptosis [10]. Therefore, iron can play a dual role as a pro- and anti-survival factor by participating in promotive/inhibitive reactions related to ROS generation and DNA synthesis. Iron chelators can isolate iron away from the cell and deprive cancer cells of their highly required source, and hence decrease viability and proliferation. They can also change the basal ROS level in cells [7]. They decrease ROS generation by sequestering iron from Fenton reaction. On the other hand, iron chelators have been shown to increase ROS generation in different ways such as, prooxidant effects, binding cellular iron and downregulating DNA synthesis, and increasing the labile iron pool (LIP) [11]–[14]. As the basal level of ROS in cancer cells is higher than normal cells, they are more sensitive to upregulation of ROS. Such effect has been attracting attention on investigating the therapeutic effects of these chelators on cancer cells [15], [16]. Iron chelating drugs have attracted attention as potential therapeutic drugs for cancer therapy. Because they upregulate the expression of hypoxia-inducible factor 1-α (Hif1-α) and have short circulation half-life, they have tumor suppressing effects by causing iron depletion. Deferoxamine (DFO) and deferiprone (DFP) are FDA-approved iron chelators routinely used to alleviate systemic iron burden in thalassemia, and are in clinical trial for Alzheimer’s (AD) and Parkinson’s (PD) diseases [17], [18]. DFO known for its efficacy in iron binding and DFP with the ability to penetrate cells were separately used to gain a comprehensive understanding of their distinct contributions to iron homeostasis in our experimental model.

The hypoxic microenvironment in tumors contributes to chemoresistance by inducing adaptive responses and altering cellular metabolism, leading to reduced drug efficacy and treatment resistance in these aggressive cancer cells [19]. Non-physiological levels of oxygen tension in hypoxia alter cancer cell metabolism and lead to angiogenesis, epithelial-to-mesenchymal transition (EMT), upregulate proinflammatory cytokines, etc., which in turn contribute to therapy resistance [20]. Hif1-α is a subunit of DNA binding protein that forms a transcriptionally active complex to activate the hypoxia gene [21]. Hif1-α subunit contains oxygen-dependent degradation (ODD) which interacts with a family of poly1-hydroxylase enzyme (PHD), in well oxygenated condition, and leads to hydroxylation. In low levels of oxygen, however, PHD is inhibited and therefore Hif1-α is stabilized. In general, the oxygen levels in different types of hypoxic tumors range from 0.3-4.2%, where the median for the brain tumors is around 1.7% [22]–[24].

Chemoresistant is associated with increased glucose uptake in cells, supporting their enhanced energy demands and promoting cell survival under stress [8]. Glucose has a critical physiological role in cell growth and survival and modulates various signaling pathways and provides energy for cell proliferation and synthesis of macro-molecules such as proteins, nucleic acids, and lipids. Also, cancer cells have the ability to switch their metabolism into aerobic glycolysis, so called the Warburg effect, by increasing glucose uptake in order to promote proliferation and long-term survival [25]. Hypoxia notably exerts a direct influence on basal glucose uptake in mammalian cells, where cell growth and survival become highly dependent on glucose concentration [26]. This influence is magnified in a tumor where its three-dimensional (3D) structure imposes nonuniform glucose and oxygen distributions due to diffusion. In addition, studies have shown that autophagy plays dual roles in TMZ-resistant GB’s response to chemotherapy and radiotherapy [27], and therefore the modulation of autophagy pathway has been receiving attention as a new threptic approach in chemo-resistant GBs [28]. The exact mechanism of resistance is not clear and understanding mechanisms underlying drug resistance and identifying novel modalities of therapy has remained as the major focus of research in the field [29].

In this work, for the first time, we investigated metabolic vulnerabilities of TMZ-resistant glioma cells to identify potential therapeutic options for treating GB. To this aim, we utilized a hybrid approach that combines theoretical modeling with 3D tumor on chip technology to explore the possibility of enhancing chemotherapy response in glioblastoma cells using metabolic intervention. In this regard, we evaluated the antitumor effects of iron and oxygen deficiency on U251 glioblastoma, using DFO and DFP, with the aim of breaking temozolomide resistance. To this aim, we first characterized the molecular signature of TMZ-resistant *vs.* non-resistant GB in 2D and 3D (see Box 1). Moreover, several downstream variations such as viability, proliferation, autophagy, intracellular iron content, RNR mRNA, apoptosis, and ROS were monitored in TMZ-resistant *vs.* non-resistant human glioblastoma cell lines (see Box 2). Finally, an *in-vitro* tumoroid model of human-derived glioblastoma was used to recreate the physiological relevant conditions of GB microenvironment and to analyze the effect of iron depletion on the viability, growth, and invasion of non-resistant and TMZ-resistant tumoroids.

**Box 1.**
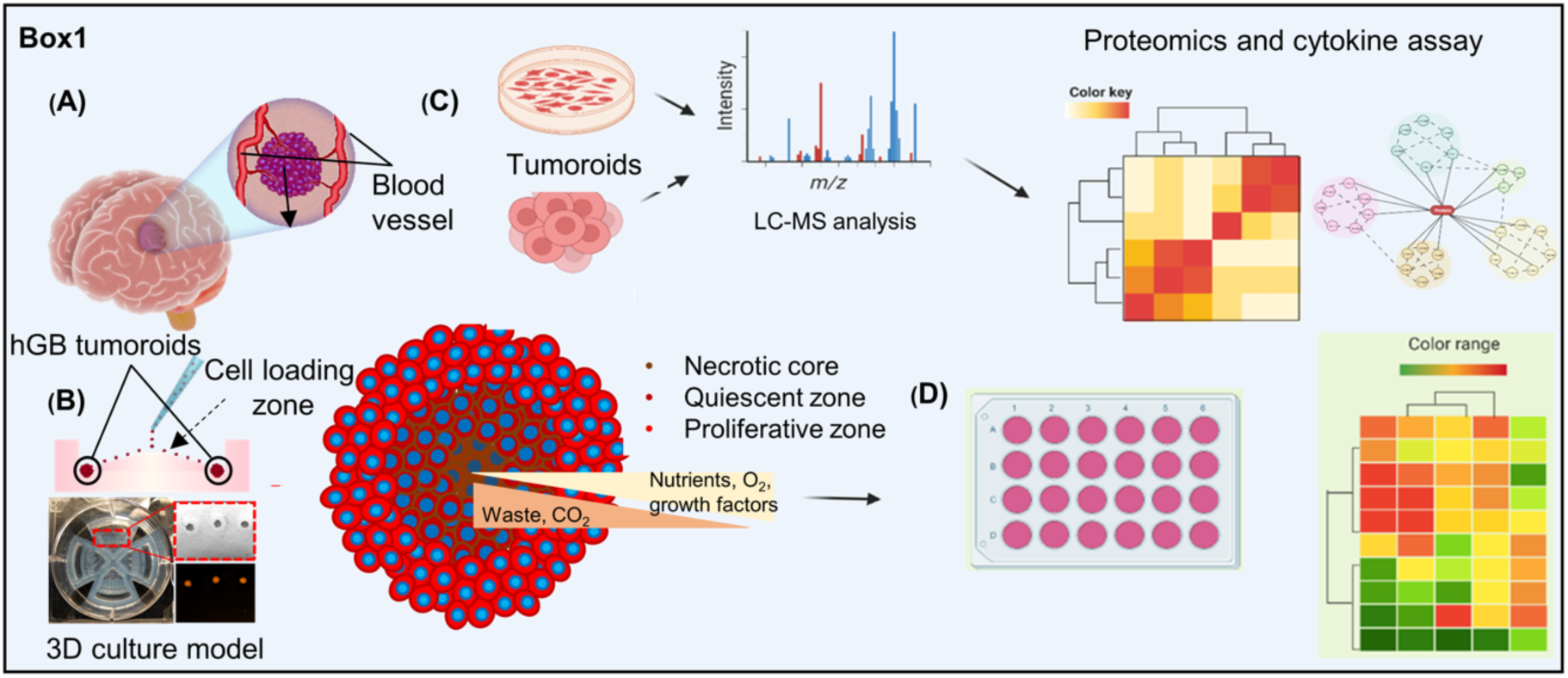
Characterization of human glioblastoma (hGB) TMZ-resistant *vs.* non-resistant. A schematic of a brain tumor and surrounding tissue. (A) Formation of non-resistant and TMZ-resistant tumoroids using EZ-Seed culture platform. (B) Tumoroids resided within a hydrogel matrix mimicking brain tumor niche in tumor microenvironment (TME). (C) workflow of the study. Steps on recapitulating resistant human glioblastoma and characterization of resistant tumoroids using proteomics, and (D) secretome analyses.

**Box 2.**
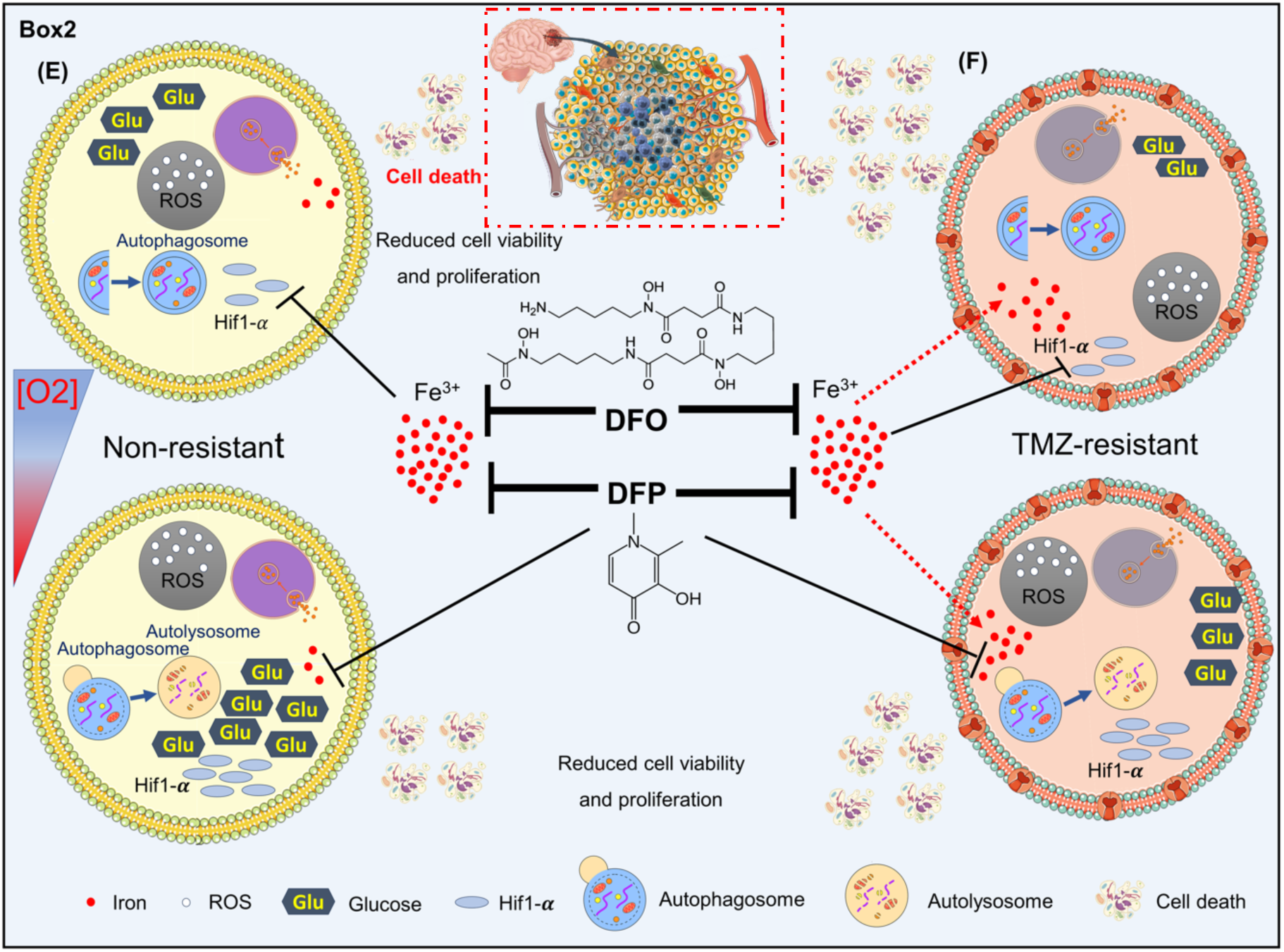
Downstream variation induced by iron and oxygen deficiency. (E) Iron deficiency induce variations in Hif1-*α* expression, ROS generation, autophagic flux, RNR, viability, proliferation, and cell death, in hGB non-resistant, and (F) TMZ-resistant cells. Graph is prepared in Biorender.

## Results

We established a TMZ-resistant glioma cell line using a pulsed-selection strategy [30]. Live/dead imaging of U251 cells after four days encapsulation in CH (Fig. S1A) and confocal imaging of actin filaments, visualized with phalloidin staining, comparing the invasion pattern of non-resistant vs. TMZ-resistant tumoroids (Fig. S1B) verified the compatibility of the hydrogel. TMZ-resistant U251 cells with a half-maximal effective concentration (EC50) of 674 µM, which was six times higher than the non-resistant cells (111 µM), were obtained (Fig. S1C). The EC50 for TMZ-resistant tumoroids was 5000 µM, whereas this value was 250 µM for non-resistant tumoroids (Fig. S1D). We evaluated the sensitivity of cells grown in 2D and 3D tumoroid cultures. To this aim, tumoroids (200 µm in diameter) were formed using EZ-Seed culture platform (Apricell Biotech, Fig. S1E) and were treated with a range of 0-5000 µM TMZ for up to four days. Both non-resistant and TMZ-resistant tumoroids displayed significantly lower chemosensitivity when compared to 2D cultures (Fig. S1F).

First, the variations of intracellular iron content, RNR mRNA, Hif1-α, autophagic flux, viability, proliferation, and ROS in response to iron depletion and oxygen tension were studied in non-resistant and TMZ-resistant cells. Next, we used an *in-vitro* tumoroids model of human-derived GB to recapitulate the physiological relevant condition of GB. Using the 3D model, we investigated the viability, growth, and invasion of non-resistant and TMZ-resistant tumoroids in response to iron depletion.

### TMZ-resistant cells acquired characteristics of mesenchymal stemness, vesicular transport, mitochondrial respiration, and mitotic quiescent

Molecular signatures of non-resistant *vs.* TMZ-resistant cells were characterized *via* comprehensive proteomics and secretome analyses. Principal component analysis was conducted to gain insights into the biological basis of acquired resistance and the impact of 3D culturing (Fig. 1A). Acquired resistance differentiated TMZ-resistant and non-resistant 2D culture with both PC1 and PC2, while 3D culturing separates TMZ-resistant tumoroid from 2D culture with high expression on PC1 and lower expression on PC2. Hierarchical clustering identified 1100 differentially regulated proteins corresponded to 3D culture and resistance characteristics (Fig. 1B). The enrichment analysis revealed that acquiring resistance was associated with the decreased expression of lysosomal and ribosomal proteins, and increased expression of genes involved in oxidative phosphorylation and vesicular transport (Fig. 1C), which underscores the reliance on oxygen and iron metabolism [31], [32]. Moreover, 3D culturing was associated with lower expression of proteins involved in RNA transport. These findings shed light on the intricate interplay between acquired resistance and cellular metabolic processes. Here, we first identify molecular signatures of 2D *vs.* 3D cultures and subsequently investigate the acquired resistance profile in TMZ-resistant vs. non-resistant models.

**Figure 1.**
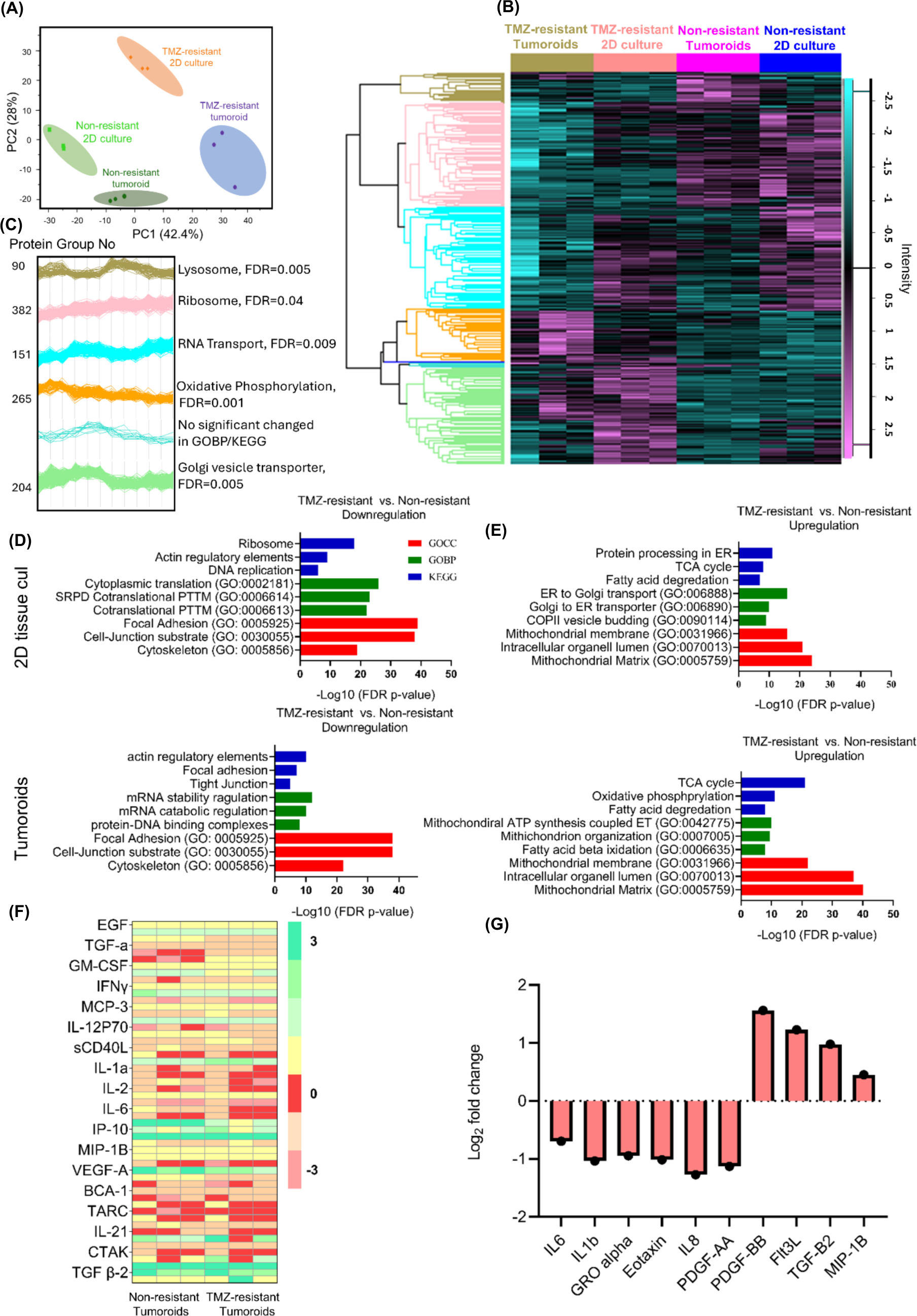
Global characterization and molecular signature of U251 non-resistant and resistant in 2D and 3D models. (A) Principal component analysis demonstrates a clear distinction between the proteomes of 2D tissue culture and 3D tumoroid in non-resistant and TMZ-resistant samples. (B) Hierarchical clustering of normalized protein concentrations. Each raw represent a distinct protein and each column represent a sample (N=3 biological independent experiment and one-way ANOVA: Benjamini–Hochberg FDR = 0.05). (C) Protein expression profiles for each cluster and the most enriched KEGG/GOBP per cluster is shown on the right side of each profile. (D&E) The three most significant GO terms and KEGG pathways that upregulated or downregulated in each group emerged from the enrichment analysis of the genes identified by student t-test. (F) Hierarchical clustering of normalized cytokine concentrations. Each raw represent a distinct cytokine and each column represent a sample (significant cytokine was extracted based on the difference in their expression values with log2 fold change ≥ 2, FDR = 0.001). (G) Cytokine profile comparing non-resistant and TMZ-resistant tumoroids, N=3 biological independent experiment.

The differentiation between 3D-cultured resistant cells and their 2D counterparts within the principal component analysis hints at a distinct metabolic shift in the 3D microenvironment, likely linked to the interplay of iron metabolism, hypoxia adaptation, and glucose utilization, underscoring the critical role of metabolic adaptations in acquired drug resistance. We performed a volcano plot analysis comparing TMZ-resistant and non-resistant cells in 2D and 3D cultures to gain insight into the differences between 3D culture and 2D monolayer cells at the protein level (Fig. S2B,C). Upregulated proteins in resistant cells were associated with decreased programmed cell death (BST2, FTL and FTH1 and ASAH1), induction of mesenchymal stemness (ALDH2 and SERPINH1), and cell cycle arrest (3IFITM, ISG15, and p53). Downregulated proteins indicate decreased glial fibrillary acidic protein (GFAP) and lysosomal activity. In this study, we observed that the stemness of cells within tumoroids was more evident when compared to cells cultured in 2D. This was because of the higher reduction in the expression of GFAP in both TMZ-resistant and non-resistant tumoroids. These findings underscore the significant impact of 3D culture conditions on cell phenotype *in-vitro*, suggesting that 3D tumoroids better represent the characteristics of cells within the tumor microenvironment (TME).

The increased expression of genes related to oxidative phosphorylation in TMZ-resistant cells may signify heightened energy production, potentially contributing to their ability to withstand the cytotoxic effects of the drug. Additionally, the lower expression of lysosomal and ribosomal proteins suggests alterations in cellular recycling mechanisms and protein synthesis, possibly aiding in resistance development.

The invasion pattern is closely linked to GBM progression, depicting the infiltration of cells to the surrounding brain tissue. The non-resistant tumoroids developed a star-shaped invasion pattern with extended protrusions into the healthy areas of the matrix, resembling diffuse gliomas. On the other hand, TMZ-resistant tumoroids exhibited a scattered tumor growth with dispersed protrusions into the surrounding matrix. A closer look at the invading cells disclosed that the non-resistant cells were elongated with clustered actin, finger-like projections into the surrounding TME, and filopodia protrusions, seen in mesenchymal cells [33], [34]. Conversely, resistant cells were rounded with bleb-like projections that are shallow and actin-rich, podosomes protrusions, seen in amoeboid cells [33], [34]. This migration plasticity confirms decreased adhesion properties in resistant cells related to mesenchymal to amoeboid transition (MAT), one of the stem cell characteristics [35]–[37].

Furthermore, Cathepsin L1 (CTSL) exhibited a significant decrease in 3D resistant cells compared to 3D non-resistant cells, indicating lower lysosomal protein expression in 3D cultures as supported by the enriched KEGG pathway in cluster 1. Protein-protein interaction networks were constructed using Network Analyst. Enrichment analysis revealed the involvement of citric acid cycle (TCA), fatty acid degradation, and mitochondrial structures (Fig. 1D&E). Notably, in 3D tumoroids, ATP synthesis and fatty acid beta-oxidation were enriched. Downregulated pathways involve actin cytoskeleton regulation, focal adhesion, and tight junctions. Protein-protein interaction networks highlighted the TCA cycle and respiratory electron transport as upregulated pathways, while translation and apoptosis are downregulated. The increased expression of genes related to oxidative phosphorylation in TMZ-resistant cells may signify heightened energy production, potentially contributing to their ability to withstand the cytotoxic effects of the drug. Additionally, the lower expression of lysosomal and ribosomal proteins suggests alterations in cellular recycling mechanisms and protein synthesis, possibly aiding in resistance development.

Actin filaments play a central role in the dynamic remodeling of the cellular cytoskeleton. In cancer cells, invasive behavior is often characterized by significant alterations in cytoskeletal architecture [38]. Confocal imaging of actin filament and compared invasion patterns of resistant and non-resistant tumoroids (Fig. S2B). The non-resistant tumoroids developed a star-shaped invasion pattern with extended protrusions into the healthy areas of the matrix, resembling diffuse gliomas. On the other hand, TMZ-resistant tumoroids exhibited a scattered tumor growth with dispersed protrusions into the surrounding matrix. A closer look at the invading cells disclosed that the non-resistant cells were elongated with clustered actin, finger-like projections into the surrounding TME, and filopodia protrusions, seen in mesenchymal cells [33], [34]. Conversely, resistant cells were rounded with bleb-like projections that are shallow and actin-rich, podosomes protrusions, seen in amoeboid cells [33], [34]. This migration plasticity confirms decreased adhesion properties in resistant cells related to mesenchymal to amoeboid transition (MAT), one of the stem cell characteristics [35]–[37]. Furthermore, Cathepsin L1 (CTSL) exhibited a significant decrease in 3D resistant cells compared to 3D non-resistant cells, indicating lower lysosomal protein expression in 3D cultures as supported by the enriched KEGG pathway in cluster 1.

To gain an insight into the effect of TME on resistance characteristics, we obtained the secretome profile of 68 cytokines from the inflammatory/invasion family, among which 18 cytokines were significantly altered (Fig. 1F). A significant increase in the level of anti-inflammatory cytokines, including Fms-like tyrosine kinase 3 ligand (Flt3L), TGF-β2, and PDGF-BB were detected in TMZ-resistant tumoroids compared with non-resistant counterparts (Fig. 1G). Further, a significant decrease in the level of pro-inflammatory cytokines such as IL-6, IL-8, IL-1β, and Gro-alpha in TMZ-resistant tumoroids were detected, compared with non-resistant cells (Fig. 1G). These findings support the decreased inflammatory reactions we observed in proteomic studies of resistant tumoroids.

Overall, acquired resistance was characterized by increased expression of genes related to DNA repair, cell survival, vesicular transport, the TCA cycle, and oxidative phosphorylation, while genes involved in cell differentiation and cellular connections, i.e., junctions and cytoskeleton, and inflammatory reactions are downregulated. The elevated reliance of TMZ-resistant cells to iron and oxygen for their metabolism and survival revealed their metabolic vulnerabilities.

### TMZ-resistant cells acquired distinctive traits in their response to iron chelators

We studied the effects of iron deficiency in 2D culture for preliminary dosage screening of these drugs. Tumors often develop hypoxic zone due to limitation in oxygen diffusion. To account for this, we considered both normoxic and hypoxic states of cells. Intracellular iron, viability, proliferation rates, and RNR mRNA level were measured in non-resistant and TMZ-resistant cells in response to 24h of treatment with DFO and DFP in the range of 10-100 μM, in normoxia and hypoxia.

DFO and DFP reduced intracellular iron content in normoxia except for DFO-treated TMZ-resistant cells (Fig. 2A-top panels). DFO reduced intracellular iron of non-resistant cells in a dose-dependent manner with 100 μM of DFO being the most effective condition (Fig. 2A-top left). Despite being a bidentate chelator compared to DFO, which is hexadentate, DFP was still able to comparably reduce intracellular iron owing to its lipophilicity.

**Figure 2.**
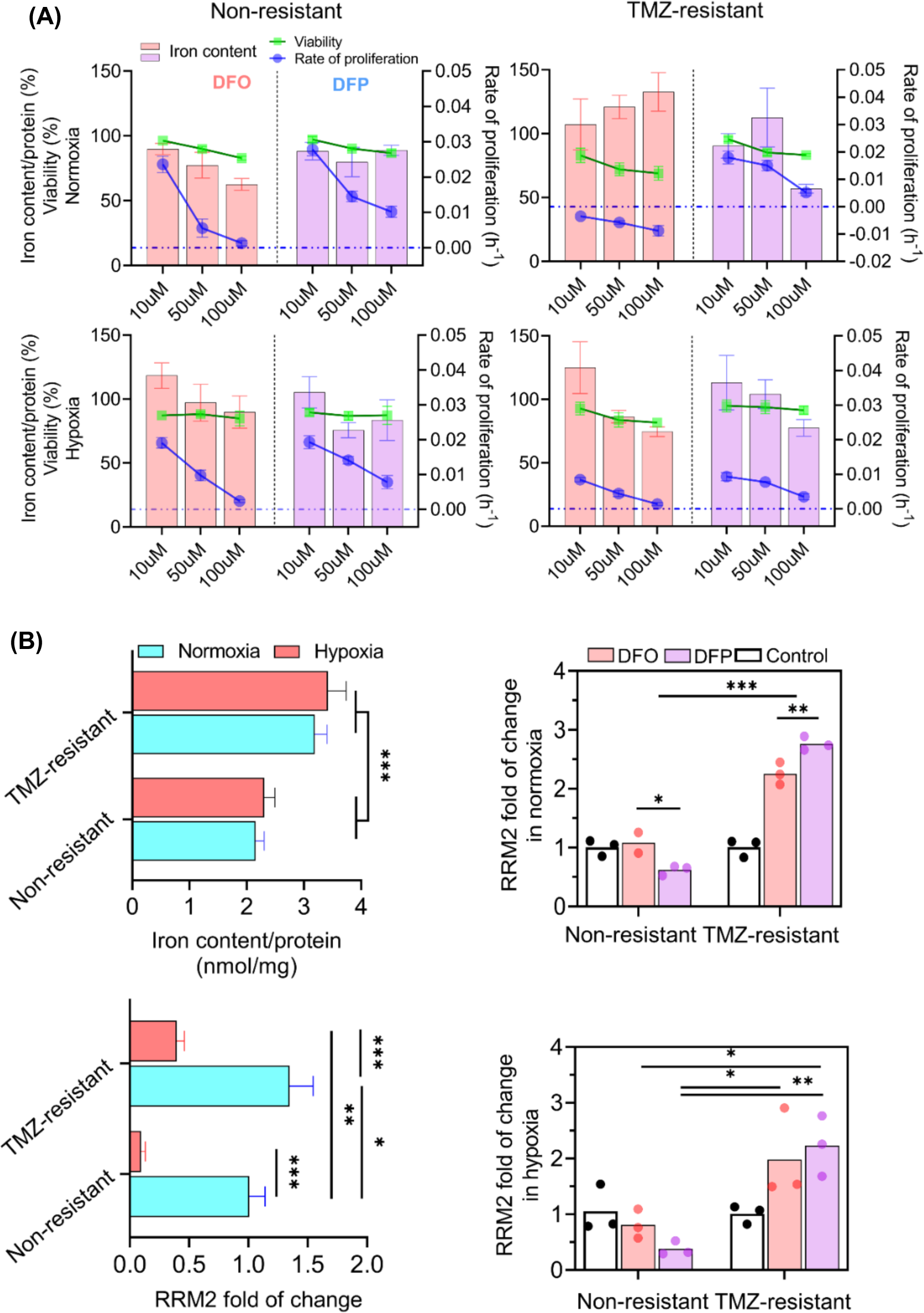
TMZ-resistant cells exhibited different pattern of variations of in viability, proliferation rate, and ribonucleotide reductase mRNA (RRM2) expression. DFO decreased intracellular iron in normoxic non-resistant cells (A-top left) but increased it in TMZ-resistant cells (A-top right). Both chelators reduced viability and proliferation, with DFO being more effective, particularly in normoxia (A-top panels). Hypoxic cells showed increased resistance to chelators (A-bottom panels). Notably, TMZ-resistant cells exhibited higher intracellular iron, potentially linked to their stem-cell-like characteristics (B-top left). TMZ-resistant cells upregulated RNR mRNA in both normoxia and hypoxia (B-bottom left). Same result was observed in TMZ-resistant cells in response to DFO and DFP in both normoxia and hypoxia (B-right panels). Both chelators changed RNR expression, with DFP having a stronger effect. In non-resistant cells, RNR mRNA decreased with chelator treatment, more significantly with DFO. N=3 biological independent experiments and statistically significant at a P-value < 0.05.

DFO relatively increased iron content in normoxic TMZ-resistant cells, likely due to a positive feedback loop in cellular uptake of transferrin (Fig. 2A-top right). Reduction in intracellular iron was also seen in hypoxic cells in response to DFO and DFP (Fig. 2A-bottom panels). In non-resistant cells, hypoxia counteracted possibly by compensating the iron reduction via upregulation of genes involved in iron transport, such as transferrin (Fig. 2A-bottom left). Such a response, however, was not observed in TMZ-resistant cells, especially in DFO treatment, suggesting a strong dependency to oxygen levels (Fig. 2A-bottom right).

Next, we investigated the possible correlation between intracellular iron level and viability and proliferation rate. DFO and DFP reduced cell viability and proliferation rates of both hypoxic and normoxic non-resistant and TMZ-resistant cells (Fig. 2A). Both non-resistant and TMZ-resistant cells were more sensitive to DFO and DFP in normoxia (Fig. 2A-top panels). Hypoxic cells were able to maintain higher viability in response to the chelators (Fig. 2A-bottom panels). The same response was observed after 72h of treatment (Fig. S4A). In general, DFO treatment was more effective in all conditions in comparison with DFP. Additionally, TMZ-resistant cells were more sensitive to the drugs in all conditions in comparison non-resistant cells. Overall, the highest reduction in viability was observed in normoxic TMZ-resistant cells (Fig. 2A-top right). Of note, it’s intriguing that both cell types displayed greater susceptibilities to DFO than to DFP, and the application of DFO to TMZ-resistant cells resulted in noticeable negative proliferation rates (Fig. 2A-top right). The findings emphasize the heightened responsiveness of normoxic TMZ-resistant cells to DFO treatment.

In contrast, hypoxic TMZ-resistant cells exhibited reduced susceptibility to DFO intervention. These findings also imply that a direct and necessary correlation between intracellular iron levels and viability and/or proliferation rate may not exist. Moreover, TMZ-resistant cells showed relatively higher intracellular iron content compared to non-resistant cells (Fig. 2B-top left), possibly due to their stem-cell like characteristics [39], [40]. Hypoxic non-resistant cells acquired higher intracellular iron. We postulate that Hif1-*α* upregulated the expression of genes involved in iron transport, such as transferrin [41].

Building on the distinct responses observed in TMZ-resistant *vs.* non-resistant cells, we measured gene expression level and regulation of RNR, the rate-limiting enzyme in the synthesis of deoxyribonucleoside triphosphates (dNTPs) for DNA synthesis and repair [42], [43]. β-subunit (R2) is a homodimer encoded by RRM2 genes and produces dNTPs for nuclear DNA replication and repair [44]. TMZ-resistant cells elevated RNR mRNA level in both normoxia and hypoxia (Fig. 2B-bottom left). Same result was observed in TMZ-resistant cells in response to DFO and DFP in both normoxia and hypoxia (Fig. 2B-right panels). DFP significantly upregulated the RNA in both conditions, compared to DFO. Unlike TMZ-resistant cells, non-resistant cells downregulated RNA level in response to the chelators with DFO being more effective. These results suggest that TMZ-resistant cells upregulate RNR mRNA production in response to RNR enzyme inhibition by the chelators. The RNA level, however, reduces in non-resistant cells due to the enzyme inhibition. In both cell types, DFP showed stronger effects due to the lipophilicity characteristic. These findings suggest a nuanced relationship between intracellular iron levels and cellular responses, whereas a correlation between dosage of chelators and viability and proliferation rate.

### TMZ-resistant cells exhibited lower Hif1-α and higher ROS accumulations compared to non-resistant cells in response to iron chelators

As hypoxia displayed noticeable interference in the cellular responses of cells to iron deficiency, we further studied the role of hypoxia by measuring glucose uptake, as an indicator of cellular metabolism, and ROS production in both non-resistant and TMZ-resistant cells under hypoxia.

Expression of Hif1-α was imaged by immunofluorescence (IF) microscope, semi-quantified using flowcytometry analysis, and confirmed with western-blotting. Hif1-α accumulation was shown in non-resistant cells exposed to 24h hypoxia and 50 μM DFO (Fig. 3A-left panels), and 50 μM DFP and combination of hypoxia and either DFO or DFP (Fig. S3A). Treatments with DFO and DFP upregulated the expression of Hif1-α in both non-resistant and TMZ-resistant cells, similar to the exposure to hypoxia. The treatment with either DFO or DFP in hypoxia did not result in significant change of the expression of Hif1-α.

**Figure 3.**
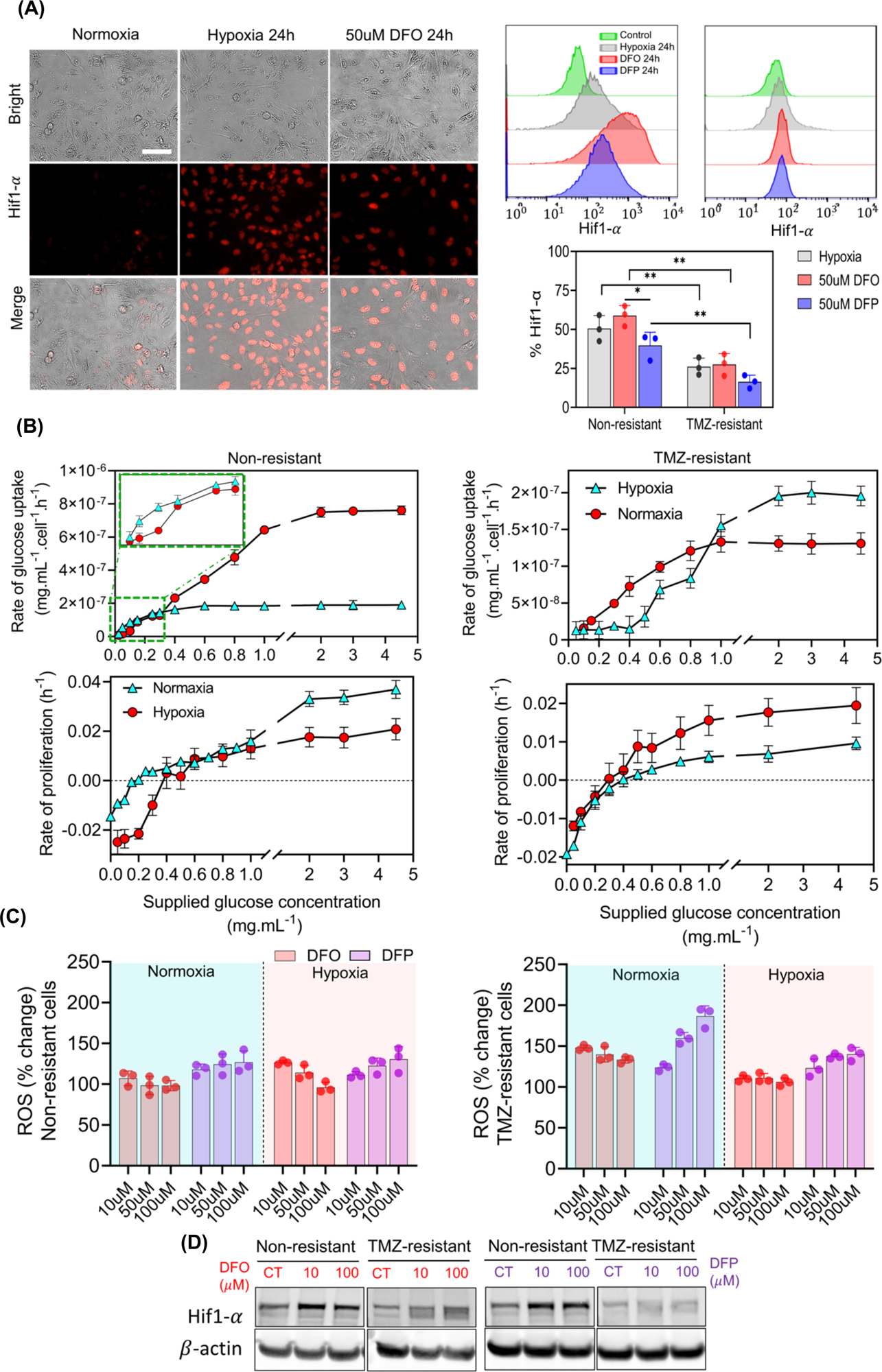
DFO and DFP upregulate Hif1-α expression and ROS generation in U251 non-resistant and TMZ-resistant cells. (A) Hif1-α expression, assessed through IF microscopy and flow cytometry (A-left panels) as well as western blotting (D) revealed upregulation in response to 24h hypoxia and 50 μM DFO and DFP treatments. Flow cytometry further quantifies Hif1-α accumulation, indicating higher levels in DFO-treated non-resistant cells compared to DFP and hypoxia, with a similar trend observed in TMZ-resistant cells (A-right panels). Glucose uptake analyses (B-top panels) demonstrate a pronounced difference between non-resistant and TMZ-resistant cells, with varying glucose concentrations affecting proliferation rates (B-bottom panels), revealing a critical threshold and different responses under hypoxic and normoxic conditions. This denotes the high sensitivity of normoxic TMZ-resistant cell to DFO. Unlike normoxic TMZ-resistant cells, hypoxic TMZ-resistant cells were less sensitive to all tested concentrations of DFO an DFP. (C) Non-monotonic dose-dependent increase of ROS in DFO-treated U251 non-resistant and TMZ-resistant cells. DFP increased ROS level monotonically in all conditions. Hypoxia reduces the effect of both DFP and DFP in TMZ-resistant cells. N=3 biological independent experiments and statistically significant at a P-value < 0.05

Western blot studies also revealed an accumulation of Hif1-α in both DFO- and DFP-treated non-resistant and TMZ-resistant cells (Fig. 3D). These results show that removing either oxygen or intracellular iron can stimulate the expression of Hif1-α. Therefore, it is important to consider the downstream signaling pathways and transcriptional changes mediated by Hif1-alpha in DFO/DFP treatment.

Additionally, the expression of Hif1-α was semi-quantified by flowcytometry analysis (Fig. 3A-right panels, Fig. 3SB). Twenty-four hours of incubation in hypoxia and 50 μM DFO and DFP treatments induced the accumulation of Hif1-α in non-resistant and TMZ-resistant cells. Cells treated with DFO exhibited a higher expression level of Hif1-α compared to both DFP treatment and hypoxia incubation in non-resistant cells. A similar trend was observed in TMZ-resistant cells. Across all treatments, non-resistant cells showed a greater accumulation of Hif1-α compared to TMZ-resistant cells.

To understand the respond of non-resistant *vs.* TMZ-resistant GB cells to nutrient perturbation, we quantified glucose uptake and proliferation rates of these cells under hypoxic and normoxic conditions. The glucose supply was in the range of 0 to 4.5 mg/mL, clinically equivalent to 450 mg/dL. Even though 450 mg/dL exceeds the normal blood glucose range of 70-144 mg/dL, it’s important to note that in GBM patients, mean glucose levels can reach as high as 459 mg/dL. [45]. In all conditions, the glucose uptake increased with glucose concentration and reached a plateau at higher glucose concentrations (Fig. 3B). This difference between the glucose uptake of non-resistant was more pronounced compared to TMZ-resistant cells (Fig. 3B-top panels). Assessing the impact of varying glucose concentrations on proliferation rate revealed a negative correlation below a critical threshold, transitioning to a positive relationship beyond this pivotal point (Fig. 3B-bottom panels). The critical threshold for cells grown under hypoxic conditions (0.5 mg/mL) was higher than those that were grown under normoxic conditions (0.2 mg/mL). However, the overall proliferation rate was higher for the cells grown under normoxic conditions. Similar trends were observed for TMZ-resistant cells. These rates were shown to be concentration dependent, that is the higher glucose concentration leads to higher rates of uptake and proliferation until the saturation point.

Hypoxic cells acquire higher glucose uptake while slower proliferation rates, and DFO-treated normoxic TMZ-resistant cells had the lowest proliferation rate, compared to all other conditions, with hypoxia counteracting the reduction. Moreover, twenty-four hours of 10 μM DFO treatment on normoxic and hypoxic non-resistant cells caused a small increase in ROS level followed by monotonic decrease in response to 50 μM and 100 μM treatments (Fig. 3C-left). 10uM, 50 μM and 100 μM of DFP treatments in these cells increased ROS level. The same nonmonotonic trend was seen in normoxic and hypoxic TMZ-resistant (Fig. 3C-right). The higher concentration of chelators, the more ROS generation was observed after 72h of treatment in all tested conditions in both DFO and DFP treatments (Fig. S4B).

### Iron chelators induced autophagy in normoxia but inhibited autophagic flux in hypoxia

GFP-RFP-LC3 adenovirus was used to monitor autophagic flux. Different sensitivity of fluorescent proteins to the lysosomal acidic environment, *i.e.,* instability of GFP in autolysosomes where mRFP was relatively stable, allowed for the characterization of the autophagic compartments. The fusion of lysosomes was visualized by imaging the loss of GFP. The number of autophagosomes was counted by co-localized mRFP/GFP (yellow puncta), while autolysosomes was counted by only red puncta.

Time-matched control cells showed double positive RFP/GFP-LC3 in cytoplasm (Fig. 4A), while both DFP and DFO treated cells expressed localized GFP-LC3 puncta indicating the accumulation of cytosolic double membrane organelles, i.e., autophagosomes (Fig. 4B). The autophagic flux was observed by imaging autolysosomes (red puncta) in cells treated with Rap, while inhibition of autophagic flux was monitored by co-localization of mRFP/GFP (yellow puncta) in Rapamycin (Rap) + Chloroquine (CQ) treated cells. It was observed that DFO and DFP treated normoxic non-resistant and TMZ-resistant cells displayed the co-localization of mRFP-GFP (yellow puncta), implying an autophagic flux inhibition. Positive control cells were treated with 750 nM Rapamycin (Rap), autophagy inducer, and 750nM Rap + 30 μM CQ, autophagic flux inhibitor. Intensity plot along puncta trajectory visualized peaks mismatch, i.e., autophagic flux, in Rap-treated cells, peaks overlap, i.e., flux inhibition, in Rap + CQ-treated cells.

**Figure 4.**
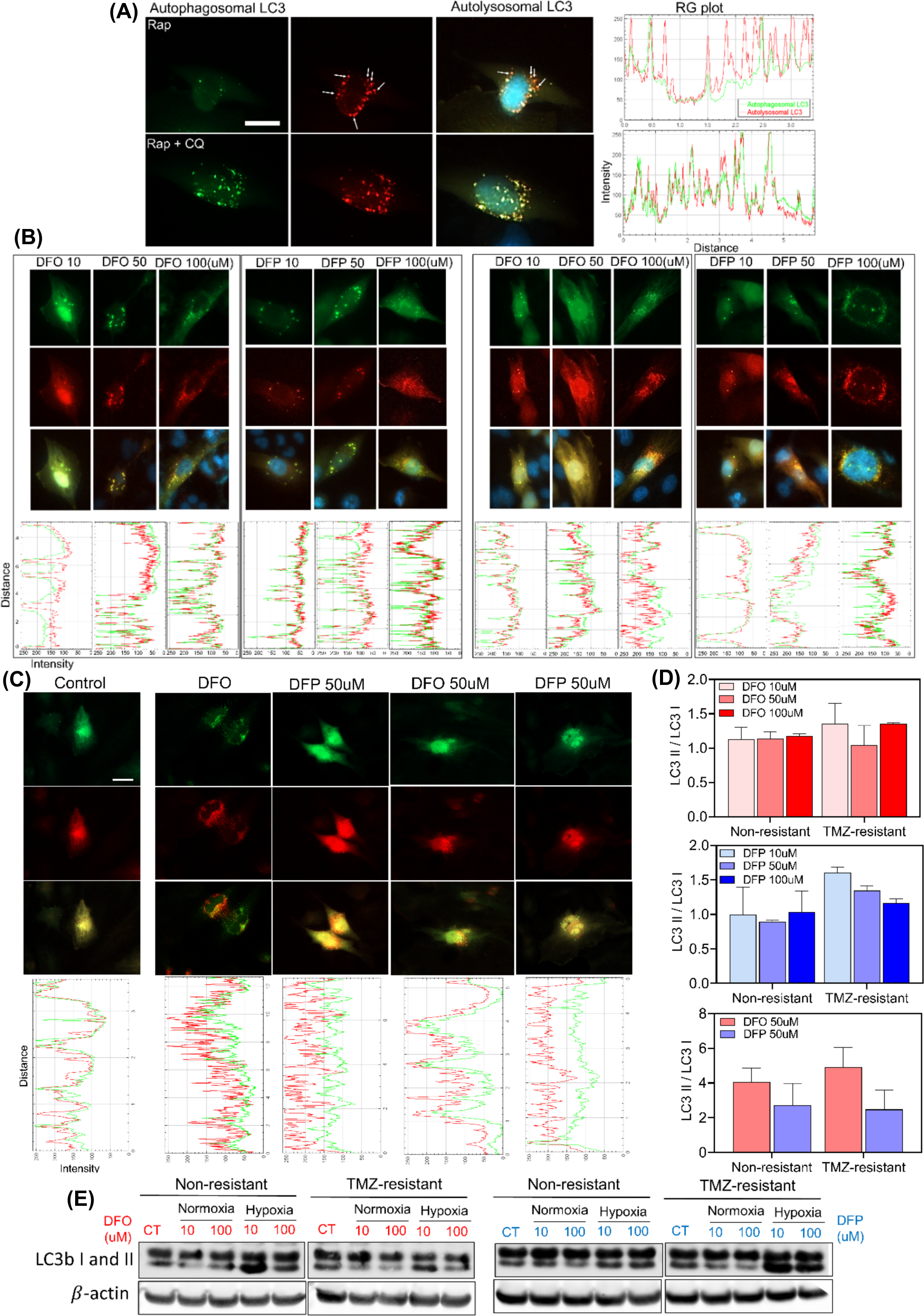
DFO and DFP induce autophagy but inhibit autophagic flux in normoxia, whereas they induce autophagy and regulate the autophagic flux in hypoxia. (A) Autophagic flux visualization using GFP-RFP-LC3 adenovirus in U251 NR and TMZ-resistant cells. Positive control cells treated with 750 μM Rap and 750 nM Rap + 30 μM CQ. The autophagic flux was observed by imaging autolysosomes (red puncta) in cells treated with Rap, while autophagy inhibition was monitored by localized mRFP/GFP (yellow puncta) in Rap + CQ treated cells. (B-left) Effects of 10uM, 50 μM and 100 μM of DFO and DFP treatments on NR cells showed co-localization of mRFP-GFP (yellow puncta) in all concentrations of DFO and DFP, which indicates the inhibition of autophagy flux. (B-right) The same treatments were performed on TMZ-resistant cells. Similar to NR cells, autophagy inhibition by DFO and DFP was observed in TMZ-resistant cells. (C) Different variation in autophagic flux was observed in hypoxic non-resistant and TMZ-resistant cells. Iron chelators induced autophagy, but regulated the flux, unlike normoxic cells. (D) The ratio of LC3II to LC3I quantified by counting red and yellow puncta confirmed the inhibition and regulation of autophagic flux in normoxia and hypoxia, respectively. (E) These results were also confirmed by WB. Scale bars are 20 *µ*m.

Next, we investigated the effects iron chelator concentration on autophagic pathway (Fig. 4B-left). A concentration dependent co-localization of mRFP-GFP (yellow puncta) was observed in all concentrations of DFO and DFP treatments indicating the inhibition of autophagy flux, confirmed by intensity plots. Similar to non-resistant cells, autophagic flux was inhibited by DFO and DFP in TMZ-resistant cells (Fig. 4B-right). Interestingly, autophagic response analyses under hypoxic conditions revealed that none of the iron chelators inhibited the autophagic flux neither in non-resistant nor in TMZ-resistant cells (Fig. 4C).

The ratio of LC3-II to LC3-I is a valuable indicator of autophagic activity. Monitoring the ratio of LC3-II to LC3-I provides insights into the autophagic flux. This ratio was quantified, equivalent to the ration of red to yellow puncta, as the measure of autophagic flux (Fig. 4D). Higher ratios in hypoxic cells indicated the fusion of lysosome, i.e., regulation of autophagic flux. These results were also confirmed by WB (Fig. 4E). In overall, results suggest that iron deficiency induce autophagy in normoxic cells, while hypoxic cells are able to persevere the autophagic flux.

### TMZ-resistant and non-resistant cells undergo cell blebbing in response to iron chelators

Cell blebbing was observed in DFO and DFP-treated non-resistant and TMZ-resistant cells. Higher concentrations of the drugs induced a larger number of blebs, with DFO being more effective (Fig. 5A-top panels). Both non-resistant and TMZ-resistant cells underwent blebbing stage and hypoxia counteracting (Fig. 5A-bottom panels). Expression of bax and caspase3 in normoxic and hypoxic non-resistant and TMZ-resistant cells are investigated using WB (Fig. 5B) and further by densitometry analyses of the WB results (Fig. S4C). Bax expression was increased in DFO-treated normoxic and hypoxic TMZR cells. In NR cells, however, the treatment did not increase bax expression. In general, DFP treatment did not affect the bax expression in neither of cells. Neither of the chelators showed significant increase in expression of caspase3

**Figure 5.**
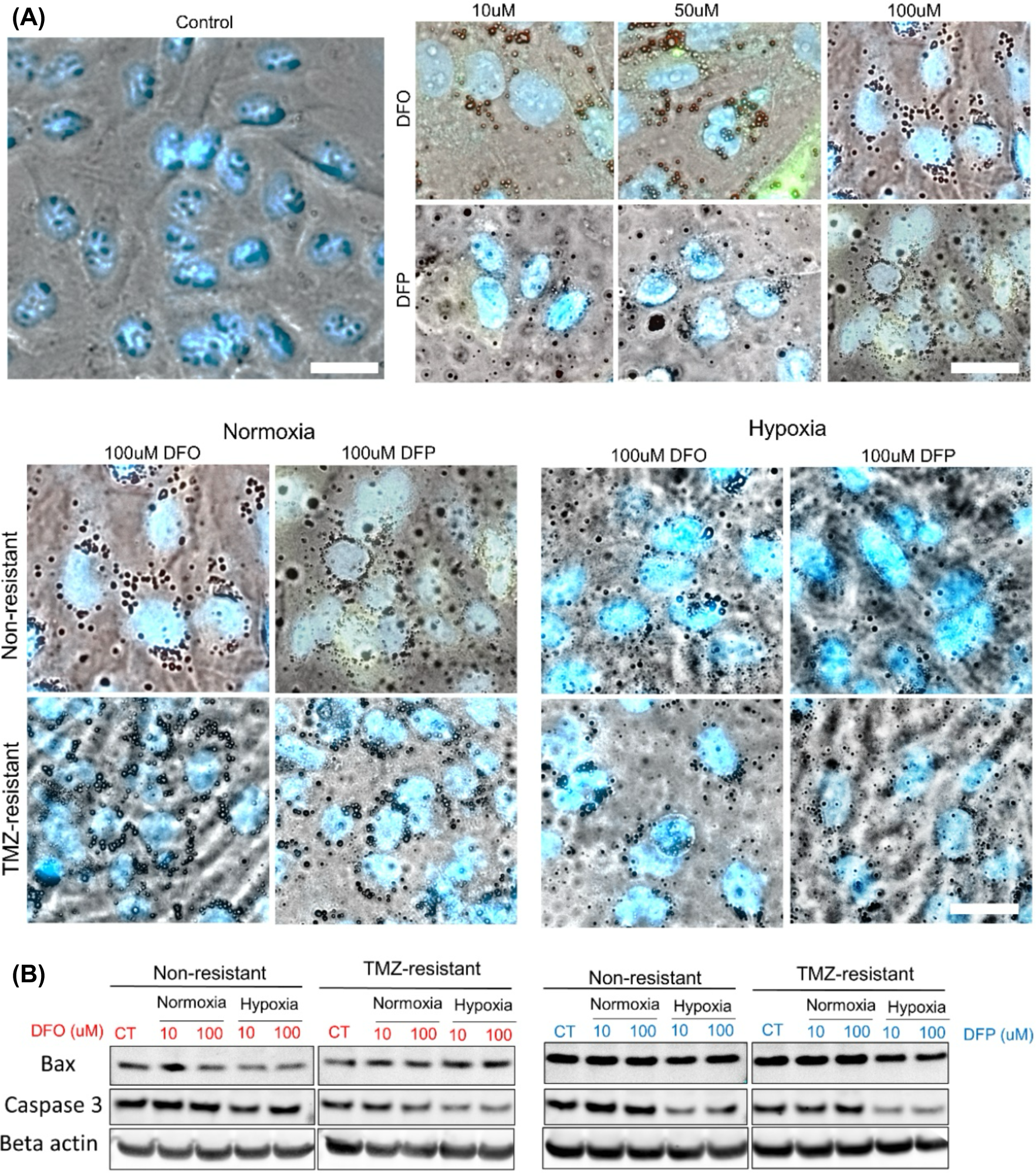
Higher expression of bax in DFO-treated TMZ-resistant cells, while hypoxia reduces the expression of caspase3. (A) Dose dependent cell blebbing was observed in normoxic non-resistant cells. Hypoxia reduced cell blebbing in both non-resistant and TMZ-resistant cells. (B) Expression of bax and caspase3 in normoxic and hypoxic non-resistant and TMZ-resistant cells in response to 10uM and 100uM of DFO and DFP. Bax expression was increased only in response DFO in normoxic and hypoxic TMZ-resistant cells. However, bax was not increased in NR cells under any treatment. Although neither of the chelators showed significant increase in expression of caspase3, it was reduced in hypoxia. Scale bars are 20*µ*m.

### Cotreatment with iron chelators TMZ induce synergic effects on hGB tumoroids

Despite tremendous efforts in developing new therapeutics for GB, most drugs have failed to improve the overall survival of patients in clinical trials. Poor clinical outcomes of developed drugs are primarily attributed to the inability of existing preclinical models to emulate the pathophysiology of GB, including CSCs population, and the 3D nature of the tumor microenvironment [46].

Using a microfluidic-integrated culture plate with U-shaped microcavities, we created an ECM-incorporated tumoroid model (Fig. 6A-left). These microcavities were connected to an open-surface microchannel, which allowed us to address each quadrant of the hydrogel with specific type and concentration of therapeutic drugs. Each quadrant of the culture plate had six microwells that acted as replicates for an experiment. We conducted an experiment to study the effect of iron-chelating drugs on viability, growth, and invasion of non-resistant and TMZ-resistant GB tumoroids. The tumoroids were screened against range of 10 to 100 μM of DFO and DFP. Live-dead imaging of cells within the tumoroid in response to the various concentrations of the DFO and DFP was illustrated (Fig. 6A-right). It indicates the ascending population of dead cells by increasing drug concentrations in both DFP and DFO treatment conditions. However, DFO induced higher toxicity on the viability of the tumoroids, confirmed by the measurement of tumoroids metabolic activity on day 1 and day 3 of the treatments (Fig. 6B). None of the treatment conditions showed toxicity levels above 50%.

**Figure 6.**
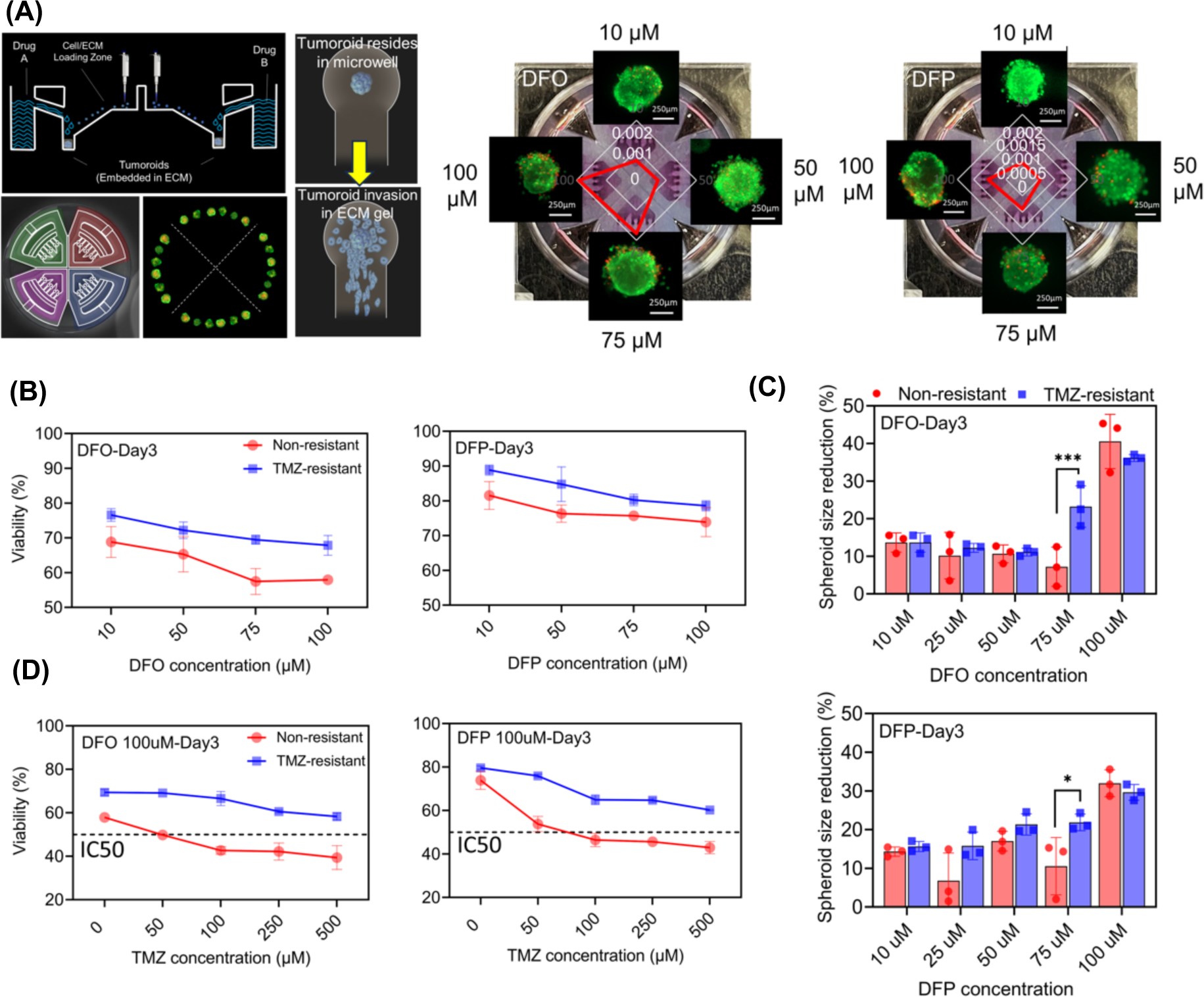
*In-vitro* hGB tumoroid in-a-well model recapitulates the physiological relevant condition in response to iron depletion. (A) Schematic side-view of the microfluidic-integrated culture plate (MiCP) for tumoroid array fabrication. Live-dead fluorescent imaging of tumoroids in each quadrant of the MiCP after treatment with varying concentrations of DFO and DFP along with semi-quantified dead cells within the tumoroids. (B) Change of viability and size reduction of tumoroids after day 1 and 3 of treatment with chelators. (C) Viability assessment of the non-resistant tumoroids treated with DFO and DFP and varying concentrations of TMZ. (D) Comprehensive heat map analysis of tumoroids viability to diffident dosages of TMZ and iron-chelator.

To determine the inhibitory effect of iron-chelator drugs on tumoroid progression, we measured tumoroid size reduction on day 1 and day 3 of the treatments. We observed a significant reduction in tumoroid size at day 3 (around 40%) after exposure to 100 μM DFO compared to lower dosages. However, the DFP treatment showed a significant size reduction effect on at 100 μM concentration on both day 1 and day 3, indicating higher sensitivity of the tumoroids to DFP treatment compared to DFO treatment.

We also studied the synergic effect of multidrug treatment on tumor progression. We exposed arrays of both non-resistant and TMZ-resistant tumoroids to varying concentrations of TMZ, ranging from 50 μM to 500 μM, combined with the lowest and highest dosages of iron chelators to sequentially expose the model to different concentrations of TMZ. Results indicated that the time of treatment has a more promising effect on reducing tumoroid viability in both non-resistant and TMZ-resistant tumoroids, as opposed to the initial concentrations of the iron chelators. Specifically, after three days of exposing non-resistant tumoroids to DFO at dosages of 10 and 100 μM, the half maximal inhibitory concentration (IC50) of the tumoroids was determined to be at a concentration of 100 μM of TMZ (Fig. 6C-top left). However, no IC50 was observed in the toxicity curve of the TMZ-resistant cells after exposure to the same co-treatment condition (Fig. 6C-top right). Sequential treatment of non-resistant tumoroids with DFP and TMZ showed the IC50 dosage of 500 μM TMZ after 3 days exposure to DFP (Fig. 6C-bottom left). Similar to the DFO behavior of the TMZ-resistant tumoroids, no IC50 reported for their sequential co-treatment with DFP and TMZ (Fig. 6C-bottom right). Through cell toxicity analysis utilizing a heat map of multidrug treatment on non-resistant tumoroids, it has been demonstrated that combining DFO and TMZ at concentrations of 100 μM and 500 μM, respectively, had a synergistic effect on the toxicity of these tumoroids (Fig. 6D-left). Neither DFO nor TMZ alone displayed any significant level of toxicity. This combination resulted in a reported viability of around 40%. However, the same treatment applied to TMZ-resistant tumoroids showed 60% viability within the model (Fig. 6D-right).

### Iron chelators significantly reduced the invasiveness of tumoroids

We examined the invasive behavior of glioblastomas (GBs) by monitoring tumoroids encapsulated in collagen, a key component of the extracellular matrix (ECM) in the brain. We measured the invasion length using live-dead staining in-a-well, fluorescent microscopy, and image analysis on day six of treatment with iron-chelator and a combination of TMZ and iron-chelator drugs.

Tumoroids enclosed in the ECM hydrogel were able to penetrate through an array of microchannels linked to the microwells, in various lengths, after exposure to single and combined therapies. We administered a 100 μM dose of TMZ following treatment the tumoroid with 10 μM and 100 μM dosages of both DFO and DFP. The extent of the invaded non-resistant and TMZ-resistant tumoroids was quantified (Fig. 7).

**Figure 7.**
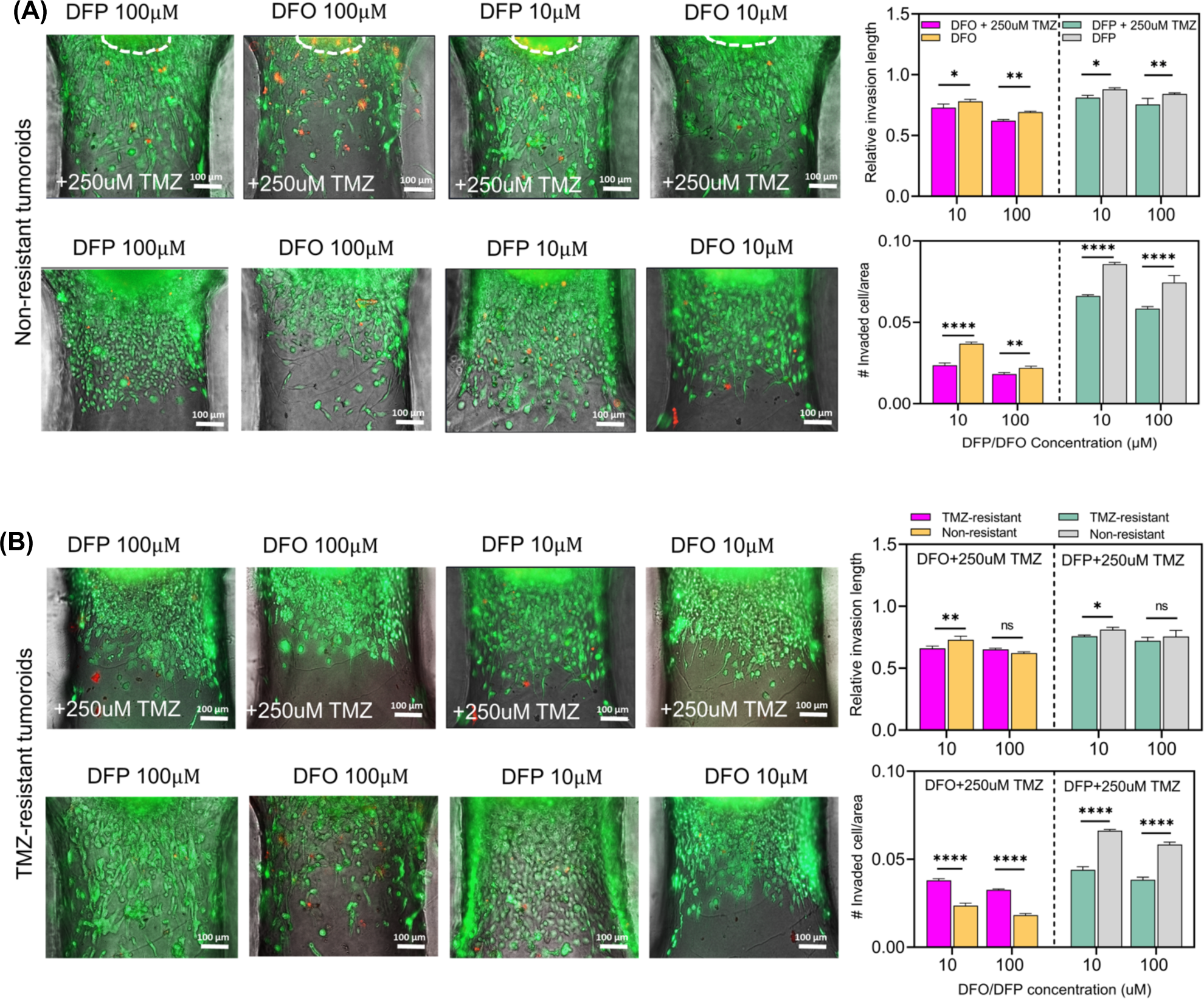
Tumoroids-in-a-well model enables invasion analysis of the ECM enclosed tumor models in response to the different combination treatment conditions. (A) Fluorescent images of the invaded non-resistant tumor cells from the tumoroid with the ECM matrix in response to different combined and single treatment conditions. (B) Fluorescent images of the invaded resistant tumor cells from the tumoroid with the ECM matrix in response to different combined and single treatment conditions. Invasion behavior of the tumoroids within the matrix was quantified through measurement of the relative invasion length of them compared to the primary tumoroid size and number of invaded cells per certain area.

These results indicated that the combined therapy had a significantly lower relative invasion length of the tumoroids in comparison to the single iron-chelator therapy for DFO. However, no significant difference was observed for the DFP treatment condition. The number of cells that invaded through the microchannel in each treatment condition was measured by counting them within a specific area. Results showed that the combination therapy reduced the lowest invasion density of tumor cells, which was consistent with previous findings. However, there was no significant difference in the number of invaded cells between 10 and 100 μM of DFO in the combination therapy. When the TMZ-resistant tumoroids were treated under the same conditions, there was no significant difference in the length of invasion between the tumoroids at different DFO dosages during the co-treatment experiment. However, in contrast to the non-resistant tumoroids, the resistant tumoroids showed a significantly reduced relative invasion length during combination therapy compared to the single DFO and DFP treatments. Furthermore, there was no discernible distinction in the density of the infiltrated cells when comparing the single and combination treatments for both 10 and 100 μM concentrations of DFO. However, the combination of DFP and TMZ treatment demonstrated a significant reduction in the density of the infiltrated cells through the ECM. This is a promising outcome towards inhibiting the diffusion of glioblastoma through brain tissue.

### Iron chelators significantly reduced the viability of recurrent GB in patient-derived cells

The viability of two types of patient-derived GB cells in response to single and co-treatment with TMZ and DFO/DFP is depicted in Fig. 8. The first cell type was derived from newly diagnosed GB tumors, labeled by BT-48, and the second type was recurrent GB cells treated by radiotherapy + concurrent adjuvant TMZ chemotherapy, labeled by BT-147, derived from a male, age 55. The results indicate that while no IC50 value was observed for either of the drugs in the single treatment regime, several IC50 values were associated with the combination in co-treatment.

**Figure 8.**
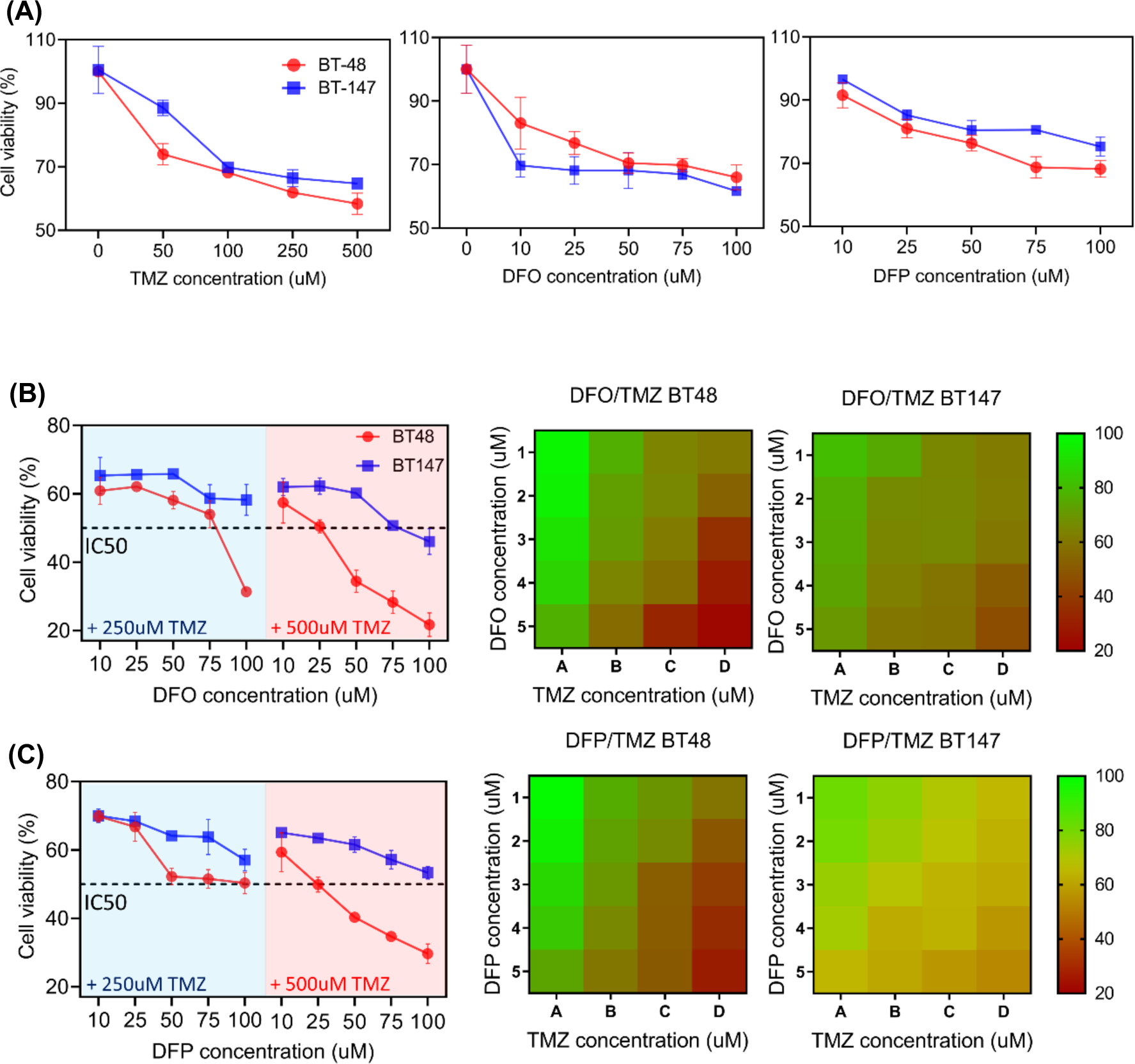
Co-treatment of glioblastoma patient-derived tumoroid with TMZ and iron chelators induced synergic effect. (A) Change of viability in response to single treatment of (left to right) TMZ, DFO, and DFP on Day 3. (B) The IC50 values were associated to the combination of 250 μM TMZ + 100 μM DFO and 500 μM TMZ + 50,75,100 μM DFO in non-resistant tumoroids, and only 500 μM TMZ + 100 μM DFO in recurrent tumoroids. (C) Additionally, the IC50 values were obtained for the combination of 500 μM TMZ + 25, 50, 75, and 100 μM DFP in non-resistant tumoroids.

In particular, TMZ-treated recurrent GB cells exhibited less sensitivity to TMZ but increased sensitivity to DFO and DFP compared to new GB cells (Fig. 8A). The IC50 values for co-treatment of new GB cells were 250 μM TMZ + 100 μM DFO and 500 μM TMZ + 50, 75, 100 μM DFO, while for recurrent GB cells, it was 500 μM TMZ + 100 μM DFO (Fig. 8B).

Additionally, IC50 values were obtained for the combination of 500 μM TMZ + 25, 50, 75, and 100 μM DFP in recurrent GB cells (Fig. 8C). These findings demonstrate that the proposed combination therapy is not only significantly effective for new GB cells but also improves efficacy in recurrent GB cells.

## Discussion

Here, we explored the potential applications of DFO and DFP in cancer treatment, particularly in relation to the role of iron in cancer cell growth and proliferation of TMZ-resistant cells. These chelators are capable of inhibiting the growth of cancer cells not only through the inhibition of DNA synthesis but also via targeting molecules involved in cell cycle control, angiogenesis and metastasis suppression [47]. Both chelators possess a sufficiently high selectivity to remove iron without, not interfering with homeostatic levels of other metals such as copper and zinc [47]. Therefore, they could serve as potential cancer therapeutics.

We observed that Iron chelators reduced intracellular iron content in a dose dependent manner in hypoxic and normoxic non-resistant and TMZ-resistant cells. However, low doses of iron chelators cannot stop TMZ-resistant cells from acquiring iron. TMZ-resistant cells acquired more iron in control condition, a stem-cell like behavior observed in GB cells. We demonstrated that the reduction of intracellular iron significantly decreased the viability and proliferation rate of cells, which were more significant in DFO-treated TMZ-resistant cells. Hypoxic cells, on the other hand, were less affected by both drugs as they are less dependent on mitochondrial metabolism and strongly rely on glycolysis. In addition, cell proliferation requires a high level of dNTPs in cancer cells. Iron chelators decrease the iron level, targeting the di-iron center of R2, hence they decreased RNR activity resulting in rate of cell proliferation. We observed that ribonucleotide reductase mRNA (RRM2) was decreased in response to DFO and DFP in non-resistant cells while TMZ-resistant cells showed elevated amount of RRM2.

Moreover, hypoxia played a counteracting role in variations of cellular responses. Significant upregulation of Hif1-α was seen in iron chelated cells and was compared with hypoxia. Similar to hypoxia, iron reduction induces Hif1-α accumulation via PHD activation. However, iron chelators reduce iron content which is normally upregulated in hypoxia. Additionally, the variation in autophagic pathway was an indicator of the important interference of iron chelators on autophagy. The interplay between autophagy and apoptotic pathways, that may either induce or inhibit apoptosis, in linked to the cellular model [48]– [50]. We observed that the autophagic flux induced by DFO and DFP shows same localization pattern, using GFP-RFP-LC3 adenovirus, and immunoblotting, in normoxic non-resistant and TMZ-resistant cells. Both drugs induced autophagy but inhibited the autophagic flux. Hypoxia on the other hand counteracted the flux inhibition.

Iron and ROS can have a dual effect in cancer cell metabolism. Even though they are shown to have a key role in cellular homeostasis, studies have shown they are important contributors to carcinogenesis. Iron can gain and lose electrons in the electron transport process on mitochondria. Controlled level of iron in healthy cells can help the normal growth and protect the cell. However, iron has an important role in the generation of ROS [10]. The reaction of iron with hydrogen peroxide yields hydroxy radical which causes DNA damage (Fenton reaction). ROS are reactive chemical groups generated due to an incomplete reduction of oxygen. Although they can damage cellular defense mechanisms, enzymes, amino acids and DNA [51], they can target kinases, cell cycle regulators and redox-sensitive transcription factors to help the regulation of cellular processes [52], [53]. Higher level of iron in cancer cells promote their growth and help them meet their need of essential factors for proliferation. Also, cancer cells have relatively higher basal level of ROS compared to normal cells and hence they are more sensitive to elevation of ROS as it leads to apoptosis and/or necrosis. Therefore, iron depletion from cellular metabolism or raising the level of ROS can be considered as therapeutic approaches in cancer treatment. Here, we observed that both chelators increased the level of ROS over 72h of treatment.

These results imply a complicated interplay between intracellular iron level and cellular response to the iron chelators, suggesting the interference of other signaling pathways in reducing the viability and proliferation rates of cells. Additionally, the combination therapy seems disrupting the pathways involved in TMZ resistance, therefore altering the invasiveness of these cells. Further studies are needed to elucidate the underlying mechanisms responsible for the observed decrease in invasion length in TMZ-resistant cells during combination therapy. Understanding mechanisms underlying drug resistance and identifying novel modalities of therapy has remained as the major focus of research in the field [29].

## Conclusion

Iron and oxygen are important cellular source for metabolism, growth and survival of cancer cells. Their roles in cancer therapeutics are complex and multifaceted. Dysregulation of iron metabolism in cancer cells, especially in TMZ-resistant GB, leads to increased iron uptake. This offers a potential approach in cancer therapy. On the other hand, hypoxia presents challenges in cancer treatment. Cancer cells in hypoxic regions are more resistant with altered drug metabolism. In this work, we studied the antitumor effects of iron deficiency, using iron-chelators DFO and DFP, on non-resistant *vs.* TMZ-resistant in 2D and 3D models of hGB with the aim of breaking temozolomide resistance in these cells. To better realize the mechanisms that lead to iron chelation-induced cell death, we monitored several downstream variations. We observed iron chelators reduced viability and proliferation, up-regulated Hif1-α expression, increased ROS generation, and induced autophagy. We observed higher expression of bax in DFO-treated TMZ-resistant cells, while hypoxia reducing the expression of caspase3. Co-treatment of hGB tumoroids with iron chelators showed potential synergic effects. The IC50 values were obtained for several combination of TMZ + DFO/DFP for both GB cell lines and patient-derived recurrence GB. The combined therapy also induced a significant reduction in the size and the relative invasion length of the tumoroids. Results of this study suggest that DFO is a promising drug candidate to improve therapeutic treatments of chemo-resistant cells. It should be noted that the roles of iron and oxygen in cancer therapeutics are complex, and their manipulation offers potential avenues to improve the efficacy of cancer treatments. However, cancer is a heterogeneous disease, and therapies targeting iron and oxygen pathways may be more effective in specific cancer types or stages. Therefore, the intricacies of these processes necessitate more profound research on their potential for novel cancer treatments.

## Materials and Methods

### Drugs, reagents, and equipment

Hypoxic incubator chamber (Cat# 27310) and a single flow meter (Cat# 27311) were purchased from Stemcell Technologies. Pre-mixed gas of 2% O_2_, 5% CO_2_ and 93% N_2_ (part # NI CD5O2C-Q) was provided by Linde Canada, Inc. Glucose oxidase (GOD, CAS# 9001-37-0), peroxidase (POD, CAS# 9003-99-0), dextrose (CAS# 50-99-7), temozolomide (TMZ, CAS# 85622-93-1), deferoxamine (DFP, CAS# 30652-11-0), deferiprone (DFO, CAS# 138-14-7), 2’, 7’-dichlorodihydrofluorescein diacetate (DCFH-DA, CAS# 4091-99-0), trypan blue (CAS# 72-57-1), presto blue (CAS# A13262) and trypsin-EDTA solution (P# T4049) were purchased from Sigma Aldrich.

Potassium iodide (CAS# 7681-11-0) from Caledon laboratory, Citric acid monohydrate (CAS# 5949-29-1) and sodium citrate (CAS# 6132-04-3) from Bio basic Canada Inc. RFP-GFP tagged adenovirus was donated by Ghavami’s lab.

Human glioblastoma cell line (hGB U251) was purchased from ATTCC (Masassas, VA, USA). Dulbecco’s Modification of Eagle’s Medium, high glucose, L-glutamine (DMEM, Cat# 11965092), DMEM, no glucose, no glutamine, no phenol red (Cat# A1443001), Fetal Bovine Serum (FBS, Cat# 10437036), penicillin-streptomycin (P/S Cat# 15140122), Hif1-α Monoclonal antibody (Mgc3) PE-conjugated (Cat# 12-7528-82), mouse IgG1 kappa isotype control PE (Cat# 12-4714-41), protease inhibitor cocktail (Cat# 87785), pierce BCA protein assay kit (Cat# 23227) and page ruler protein ladder (Cat# 26620) were purchased from Thermofisher.

NuPage 4-12% Bis-Tris gels, 1.0mm, 10-well (Cat# NP0321PK2), NuPage 10X Sample Reducing Agent (Cat# NP0009) and NuPage 4X LDS Buffer (Cat# NP0008) were purchased from Invitrogen. Intercept (PBS) blocking buffer (Cat# 927-70001) was purchased form Licor. Rabbit anti LC3B (Cat# 100-2220) was bought from Novus. Rabbit anti BAX (Cat# 92772), rabbit anti caspase3 (9662), mouse anti Hif1-α (Cat# 79233), rabbit anti beta actin (Cat# 4967), anti-rabbit IgG (H+L)-DyLight 800 (Cat# 5151) and anti-mouse IgG (H+L)-DyLight 800 (Cat# 5257) were purchased from CellSignaling. Nitrocellulose membrane was purchased form Pall (product ID# 66485).

Semi-dry Transfer Buffer was mad in house with mixing 11.64g of 48mM Tris (Cat# BDH4500, VWR), 5.86g of 39mM glycine (Cat# BP38150, ICN), 7.5mL of 0.037% SDS (Cat# L5750, Sigma) and 20% methanol (Cat# MX0485-3, EMD) with 2L H2O.

### Glucose assay

2×104 cells were cultured in 24 well plates with DMEM supplemented with 10% (v/v) Fetal Bovine Serum (FBS) and 1% (v/v) Penicillin/Streptomycin, incubated at 37 °C in 5% CO_2_ and 95% air. When cells were 50-60% confluent, the media of wells were removed and media with desired concentrations of glucose, 0.025 to 1 mg/mL, were added to each well. The rate of change in the concentration of glucose in each well were monitored over time (10h) using glucose oxidase assay with colorimetric method, following the manufacturer’s protocol. In this method, the glucose oxidase enzyme (GOD) catalyzes the oxidation of glucose to hydrogen peroxide (H_2_O_2_) and peroxidase (POD) reaction was then used to calorimetrically visualizing the formed H_2_O_2_ [30]. Also, the number of cells in each well was counted to normalize the uptake rate. It was seen that the uptake rate is constant above a threshold and below that the rate becomes dependent on the concentration of glucose. To measure the proliferation rate, number of cells before adding desired concentrations and after 24 and 48 hours of that were counted for each concentration using Trypan blue assay. To measure the rates of glucose uptake and proliferation of hypoxic cells, cultured cells were transformed into a hypoxic chamber (Stemcell Technologies; Cat# 27310) filled with a pre-mixed gas of 2% O_2_, 5% CO_2_ and 93% N_2_. After 24h the same glucose measurement and Trypan blue assays used to measure the rates of glucose uptake and proliferation.

### Rate of proliferation

After the indicated treatment times, cells were trypsinized, dissociated and centrifuged in 300g for 5min. The pallet was resuspended, and 0.4% trypan blue solution was mixed in 9:1. 10 μL of the solution was added to the hemocytometer, and number of live/dead cells was counted using immunofluorescence (IF) microscope.

### Immunofluorescence imaging

Accumulation of Hif1-α was imaged using Hif1-α alpha antibody. U251 non-resistant and TMZ-resistant cells were grown on coverslips (5×10^4^ cells per 35 mm) and treated with DFO and/or placed in hypoxic chamber. After 24h, cells were fixed using 90% methanol for 15 min in -20 °C and stained with Hif1-α antibody according to the company’s protocol. After 2h of incubation at room temperature, cells were imaged using IF microscope.

Flowcytometry was used for semi-quantification analysis. To prepare the cells for flowcytometry, cells were washed in PBS, trypsinized with trypsin-EDTA, resuspended in culture media, centrifuged at 400g for 3min, the pallet was resuspended in 90% methanol and incubated in -20 °C for 15 min. All the washing and dissociation steps were done on ice in presence of 50 μM DFO in order to inhibit Hif1-α degradation. Methanol was then neutralized with excess amount of PBS and cells were washed and centrifuged. Cell pallet was resuspended in PBS and a Hif1-α antibody was added according to the company’s protocol. To remove non-specific interactions, isotype antibody was used as negative control of the experiment.

### Cell lines and culture

Human glioblastoma multiform cells (GB U251) were cultured in DMEM with 10% (v/v) FBS and 1% (v/v) PS and maintained in 5% CO_2_ and 95% air at 37 °C in humidified standard cell culture incubator. Culture medium was replaced every other day. Monolayer cells were trypsinized, centrifuged at 300g for 5min, supernatant was removed, and the pallet was resuspended in fresh media. To count the cells, 0.4% trypan blue solution were mixed in 9:1 and 10 µL of the solution was added to the hemocytometer. Number of live/dead cells was counted using microscope.

### 2D culture

For all experiments, 10 mL of 5 x 10^5^ cell mL-1 of U251-mKate (NR) and TMZ-resistant U251-mKate cells were seeded in 100 mm Petri dishes and were nourished with TMZ-free and TMZ 250 μM-containing complete high glucose DMEM media, respectively. Cells were incubated at 37 °C and 7.5% CO_2_ in a humidified incubator for four days. The cells were scraped and washed using PBS. Cell pellets were kept in a PBS buffer containing a 1:75 ratio of phosphatase inhibitor cocktail (Sigma, Cat#P5726) and protease inhibitor cocktail (Sigma, Cat#P8340) at -80°C until further analysis.

### Proteomics Data Analysis

The mass spectrometry proteomics data have been deposited to the ProteomeXchange Consortium via the PRIDE [54] partner repository with the dataset identifier PXD037753. Raw files were created using the XCalibur 4.2.28.14 (Thermo Scientific) software. Thermo Raw files were uploaded to MaxQuant (V.1.6.2.10), and MS/MS spectra were searched against the Uniprot (Swiss-Prot) protein sequence database (Homo Sapiens, release version of May 2021) using Andromeda, with an FDR for identification set at 0.1. The MaxLFQ algorithm quantified proteins. Database search parameters were as follows: protein N-term acetylation and deamidation, methionine oxidation as variable modifications, and cysteine carbamidomethylation as fixed modification. As a result, 4122 identified proteins were uploaded into Perseus (V.1.6.2.2). Data were filtered to remove proteins identified in the reverse sequences with only modified peptides considered common lab contaminants and annotated, resulting in 3879 proteins. For the Principal Component Analysis (PCA) and Hierarchical Clustering Analysis (HCA), proteins with the non-statistical difference between sample groups, unrelated to TMZ resistance, were removed following an ANOVA analysis. The expression values of the 1100 identified proteins were normalized using Z-score. In PCA, the normalized values were plotted, considering a Benjamini-Hochberg threshold of 0.05, using the built-in tool of Perseus. In parallel, according to the Euclidean distance, HCA was performed to evaluate the relative similarities between the different groups.

The data were filtered, considering a minimum of 70% valid values in each group. The non-normalized expression values of 2024 identified proteins were log2 transformed, and missing values were replaced with the value of -1. Differentially expressed proteins between sample categories were then determined using a two-tailed t-test with 250 randomizations. The FDR was set to 0.05 and the S0 to 0.1. The data was then plotted as volcano plots. Data were exported and uploaded to JMP 16.0.0 for visualization without further processing. Upregulated proteins are shown in red, whereas downregulated are shown in blue.

Proteomics data were retrieved from the ProteomeXchange Consortium/PRIDE repository (PXD017952) for comparative analysis with human primary and recurrent tumors and analyzed in MaxQuant (V.1.6.8.0), considering an FDR set at 1% on the PSM, peptide and protein level. Spectra were compared to the human protein sequences of Uniprot (Homo Sapiens, release version of May 2021) and proteins quantified by the MaxLFQ algorithm with a minimum ratio count of two unique or razor peptides. Database search parameters were as described above. For MaxQuant, the mass tolerance of precursor mass was 20 ppm, while fragment ion mass tolerance was 0.15 Da. The minimal peptide length was defined as 7 amino acids; at most, two missed cleavages were allowed. A single matrix containing the expression of common proteins in resistant and non-resistant cells/spheroids and primary and recurrent tumors (1191 proteins) was built and uploaded in Perseus (V.1.6.2.2). As mentioned, potential contaminants, reverse identification, and proteins only identified by site were removed. Samples were grouped and filtered to contain a 70% minimum of valid values per group (1007 proteins). Non-normalized expression values were log2 transformed and missing values imputed according to a normal distribution (width 0.3, downshift 1.8). Log2 values were then Z-scored and hierarchically clustered as aforementioned. For visualization purposes, the normalized data were exported and uploaded to MetaboAnalyst 5.0 without further processing. The heatmap was then manually curated to remove clusters that do not consider proteins involved in gaining resistance to TMZ, that is, proteins with a non-significant difference between groups.

Volcano plot analysis was performed comparing TMZ-resistant and non-resistant cells in 2D and 3D cultures to gain insight into the differences between 3D culture and 2D monolayer cells at the protein level. The red or blue dots indicate proteins significantly (q < 0.05) upregulated (log2 fold change ≥ 0.5) or downregulated (log2 fold change ≤ −0.5), respectively. Proteins with more than 2-fold up or downregulation were annotated. Protein-protein interaction networks were constructed using Network Analyst [55], [56]. In brief, minimum first-order interaction networks were constructed from proteins with p<0.05 using the IMEx [57] interactome. Pathway analysis was performed on upregulated and down-regulated proteins within each network using REACTOME and Gene Ontology enrichment analysis built-in Network Analyst. Gene ontology enrichment analysis was also done on the two largest nodes with altered abundance in each network (excluding first-order interactors). Proteins with increased abundance are shown in red, and decreased abundance is shown in blue.

### Establishment of TMZ-resistant cells

U251-mKate cells (developed by Dr. Marcel Bally, Experimental Therapeutics, British Columbia Cancer Agency, Vancouver, BC, Canada) were cultured in T75 tissue culture flasks in a complete media containing high glucose DMEM (Gibco; Cat #: 11965092), 10% FBS (Gibco; Cat #: 10437-036), and 1% Penicillin-streptomycin (Gibco, Cat#:15140122) and incubated at 37 °C, 7.5% CO_2_, in a humidified incubator (Thermo Fischer Scientific). We chose a pulsed-selection strategy to make a clinically relevant chemoresistant model [34]. After reaching 70-80% confluence, cells were treated with 100 µM of TMZ (Sigma-Aldrich, CAS #:85622-93-1) over three weeks, followed by four weeks of TMZ-free media for recovery. Then, cells were cultured in complete media with 250 µM TMZ for three weeks, followed by another four weeks of culture in TMZ-free media. The final population of U251-mKate cells resistant to the 250 µM TMZ was selected and cultured in a complete media without TMZ for four weeks.

### Cell viability

1 x 10^4^ non-resistant and TMZ-resistant cells per well were seeded in 96 well plate. After 48h, three concentrations (10, 50 and 100 µM) of DFO and DFP were added to each well. Cells were incubated either in culture incubator (normoxia) or hypoxic chamber. After 24h, 48h and 72h, cell media was removed from each well and 10% presto blue solution was added. Cells were incubated for 20 minutes, the supernatant was then collected and fluorescent intensity was measured at 560/590 nm ex/em.

### Autophagy flux assay

To monitor the presence of autophagosomes, autophagolysosomes and autophagy flux we monitored the localization of LC3 using GFP-RFP-LC3 adenovirus. In brief, U251 non-resistant and TMZ-resistant cells were grown on coverslips (50000 cells per 35 mm) and treated with three concentrations (10, 50 and 100 µM) of DFO and DFP. After 24h, cells were transfected (50 MOI) with GFP-RFP-LC3 adenovirus. 24h after transfection, cell medium was removed, and coverslips were rinsed in PBS. Also, cells were post-treated with 750 nM Rap and 750 nM Rap + 30 μM CQ as the positive controls for the experiment. The fusion of lysosome was visualized by imaging the particular loss of GFP. The number of autophagosomes was counted by co-localized mRFP/GFP (yellow puncta), while autolysosomes was counted by only red puncta.

### Immunoblotting

U251 non-resistant and TMZ-resistant cells were seeded on a 3cm Petri-dishes. After 48h, they were split into two groups; cells in the first group were kept in cell culture incubator and the other group was incubated in hypoxia. After 24h, three concentrations (10, 50 and 100 µM) of DFO and DFP were added to each Petridish. In order to collect the protein, after 24h and 72h of treatments, cell media were removed, cells were washed twice with cold PBS and 400uL of cold RIPA lysis buffer (with protease inhibitor) was added to each petridishes (all steps were performed on ice). Cell scraper was used to gently scrape cells from the bottom of the dish. Cell lysates were passed through pipette 20 times to form a homogeneous lysate, and then transferred to 1.5 ml microcentrifuge tube. Samples were allowed to stand for 15 minutes at 4 °C. The mixtures were then centrifuged at 14,000 g for 10 minutes at 4 °C to separate cell debris from proteins. The supernatants were then transferred for western-blotting. Protein concentration of each sample was measured with a Pierce BCA protein assay kit. In a 1.5mL eppi, reducing agent (10x) and LDS buffer (4x) were added, double-distilled water (ddH20) to lysate to a final concentration of 2500 ug/mL, and heated at 95 °C for 10 minutes. The mixture was spud down quickly and placed immediately on ice. Precast 4-12% bis-tris gel was prepared, and after setting up the apparatus the upper buffer chamber was filled with ∼200ml MES buffer and check for leaks. Samples and 8µL page ruler protein ladder was then loaded and run for 60-75 minutes at 125V. Afterwards, gel was removed and equilibrated in semi-dry transfer buffer for 5 minutes, transferred onto a nitrocellulose membrane for 45 minutes at 22V and the membrane was blocked for 60min in odyssey blocking buffer. Primary antibodies were diluted in blocking buffer and incubated overnight at 4 °C. Membrane was washed 4 x 5 minutes in PBST (PBS + 0.1% tween 20) at room temp, then rinsed with PBS. Secondary antibodies were diluted in blocking buffer and incubated for 60 minutes at room temperature. Membrane was washed 3 x 10 minutes in PBST, then rinsed with PBS. Finally, scan blot was performed with odyssey.

### Ferrozine assay

TMZ-resistant and non-resistant U251 cells were seeded on 6 or 12 well plates and cultured until confluent. Cells were then treated with media or one of three doses of deferiprone or deferoxamine (10, 50, or 100 µM) for 24 hours. After treatment, the cells were rinsed several times with PBS and stored on the plate at -20 °C until analyzed.

To determine intracellular iron, a ferrozine-based assay was utilized as per [35], [36] and Reimer, et al., with some modifications. Briefly, the cell samples were lysed directly in well plates with a sufficient volume of Cell-lytic M lysis buffer (containing 1% protease inhibitor cocktail III and 1 mM PMSF) to cover cells. Plates were placed on an Infors HT Ecotron orbital shaker (Bottmingen, Switzerland) for 30 minutes (100 RPM, 25 °C), then transferred to microcentrifuge tubes. A small aliquot of 10 µL was taken for protein analysis. The remaining volume of lysate was dried at 95 °C in a covered VWR dry block heater (Dublin, Ireland). Cell lysates were rehydrated with 75 µL of ddH2O and 75 µL 40 mM HCl. To this solution, 37.5 µL of 1.4 M HCl and 4.5 % w/v KMnO_4_ was added sequentially, then lysates were digested at 60 °C in a VWR dry block heater. Samples were cooled, then 22.5 µL of iron detecting reagent containing 6.5 mM neocuproine, 6.8 mM ferrozine, 2.5 mM ammonium acetate and 1.0 mM ascorbic acid was added. Samples were incubated at 28 °C for 30 minutes in the dark, then centrifuged at 16000 G for 5 minutes to remove any remaining cell debris. The supernatant was plated into a 96-well plate and read at 550 nm on a multilabel plate reader. To prepare a 10-point standard curve, a range of iron standards from 0.6-150 µM were prepared using serial dilution from 300 µM FeCl_3_ in 10 mM HCl. A 75 µL volume of ddH_2_O was added to 75 µL of iron standard, then these samples were treated identically to cell lysates.

A protein assay was performed on 10 µL of cell lysate using Pierce BCA Protein Assay Kit (23225, Thermofisher) and following manufacturer’s instructions. Intracellular iron content was normalized to protein content.

### ROS assay

TMZ-resistant and non-resistant U251 cells were seed into 96-well plates. After 48h, cells were categorized into two groups, hypoxic and normoxic cells. The former was kept in hypoxic chamber and the latter was maintained in cell culture incubator. After 24h, both groups were treated with three concentrations (10, 50 and 100 µM) of DFO and DFP. After 24h and 72h of treatment, cells were rinsed twice with PBS to remove any traces of media. 100 μL of 10 µM DCFH-DA were added to each well. The plate was protected from light and incubated on a plate shaker with gentle agitation for 30 minutes at 37 °C. Fluorescent intensity was measured at 488/535 nm ex/em.

### Real-time PCR

TMZ-resistant and non-resistant U251 cells were seed into 6-well plates and were kept in normoxia and hypoxia. After 24h of treatment with DFO and DFP, they were wash with ice-cold PBS and dry pellet stored at -20 ℃. mRNA extraction and DNase digestion were performed following the instruction of Qiagen’s RNeasy Mini Kit and RNase-Free DNase kit, respectively. Extracted-RNA purity was tested using Nandrop. First-strand cDNA was synthesized following Invitrogen’s SuperScript III First-Strand Synthesis System and the purity was tested by Nandrop. QuantiNov SYBR Green PCR Kit was then used to prepare the reactions for quantitative measurement of RNR.

### Tumoroid formation on-a-plate

U251 multicellular tumoroid formation was performed using the hydrogel-based microfluidic-integrated culture plate from Apricell biotechnology (3-in-1 plate) with U-shaped microwells. Hydrogel inserts were fabricated by replica molding of the 3D printed microfeatures. Inserts in 6-well plates were seeded gently with 100 µL of culture media including 2×10^5^ cells on seeding zone of the plate. The wells were incubated at 37 °C for 10 min to let the cells fill the microwells through the microchannels of the device. Afterward, 300 µL of fresh culture media was gently added inside the media reservoir of the plate to cover the cells in microwells. U251 tumoroids were monitored daily to measure their growth over the three days.

### ECM embedding and combinatorial drug testing

To mimic the tumor ECM condition of the glioblastoma, bovine fibril collagen with concertation of 4 mg/mL was used as the main component of the tumor extracellular matrix (ECM). To prepare the hydrogel solution, first, the pH and ionic concentrations of the collagen solution were adjusted by adding 10 × PBS and 0.5 N NaOH to the stock solution of collagen with a ratio of 1:1:8. Afterwards, a certain volume of culture media was immediately mixed with collagen solution to achieve the desired concertation. The final collagen solution was gently introduced into the ECM loading zone of the culture plate insert and incubated at 37 °C for 45 min to form a gel. Various concentrations of DFP and DFO (10-100 μM) were introduced in a single treatment setting to investigate the effect of iron-chelating drugs on the metabolic activity and cytotoxicity of the brain tumoroids. To this end, 4-day-old tumoroids were treated with the drug by aspirating the primary culture medium form the media reservoir followed by adding the fresh 300 μL drug-containing fresh media. In a combinational drug treatment setting, TMZ with different concentrations of 50, 100, 250 and 500 μM was added following iron-chelating drug treatment on tumoroids on-a-plate.

### Drug toxicity and Live/Dead staining

To examine the viability of the cancer cells within the tumoroids, the Live/Dead assay was conducted using 1 μM calcein AM and 4 μM ethidium homodimer-1 (Life Technologies kit) for 30 minutes at 37 °C. The entire ECM embedded GB tumoroid model was stained and imaged on the culture plate insert without the need of tumoroid removal before staining. Effect of drug doses on single and combination treatment interventions was assessed by florescent imaging and quantification of the invaded cells with the ECM. To determine the amount of drug toxicity in each concentration and day of drug application on the tumoroid cells, Presto Blue assay was used. This assay measures the metabolic activity of the live cells with the fluorescence excitation and emission wavelengths of 560 nm and 590 nm, respectively. 10% Presto Blue reagent of the total culture medium was added directly to media reservoir of the insert and covered the tumoroids during the 3h incubation. The average fluorescence intensity of the tumoroid was subtracted from the intensity of related cell-free microwells in the presence the blank sample.

### Statistical analysis

All data represent means ± SD. We used GRAPHPAD PRISM 5.0 (Graphpad, San Diego, CA, USA) to detect the statistical significance of differences using one-way or two-way ANOVA followed by Bonferroni’s post hoc test. Differences were considered as statistically significant at a P-value < 0.05. Three independent experiments were performed unless indicated otherwise.

The Materials and Methods section should provide sufficient information to allow replication of the results. Begin with a section titled Experimental Design describing the objectives and design of the study as well as prespecified components.

In addition, include a section titled Statistical Analysis at the end that fully describes the statistical methods with enough detail to enable a knowledgeable reader with access to the original data to verify the results. The values for *N*, *P*, and the specific statistical test performed for each experiment should be included in the appropriate figure legend or main text.

## Acknowledgments

We would like to acknowledge BC cancer Foundation, Natural Sciences and Engineering Research Council of Canada (NSERC) and Canada Foundation for Innovation (CFI). We also acknowledge Dr. Saeid Ghavami’s group at University of Manitoba for donating RFP-GFP tagged adenovirus.

## Author contributions

Conceptualization: MA (Meitham Amereh), AS, PW, MA (Mohsen Akbari)

Methodology: MA (Meitham Amereh), AS, SS, SL, TZ, PW, MA (Mohsen Akbari)

Investigation: MA (Meitham Amereh), AS, MS

Visualization: MA (Meitham Amereh), AS, MS

Supervision: JL, PW, MA (Mohsen Akbari)

Writing—original draft: MA (Meitham Amereh), AS

Writing—review & editing: MA (Meitham Amereh), PW, MA (Mohsen Akbari)

## Data and materials availability

Data will be made available on request.

