## Supplemental file for "Metabolic Vulnerabilities of Temozolomide-Resistant Glioblastoma Cells: Implications for Targeted Therapies and Overcoming Chemoresistance"

### Supplementary Materials

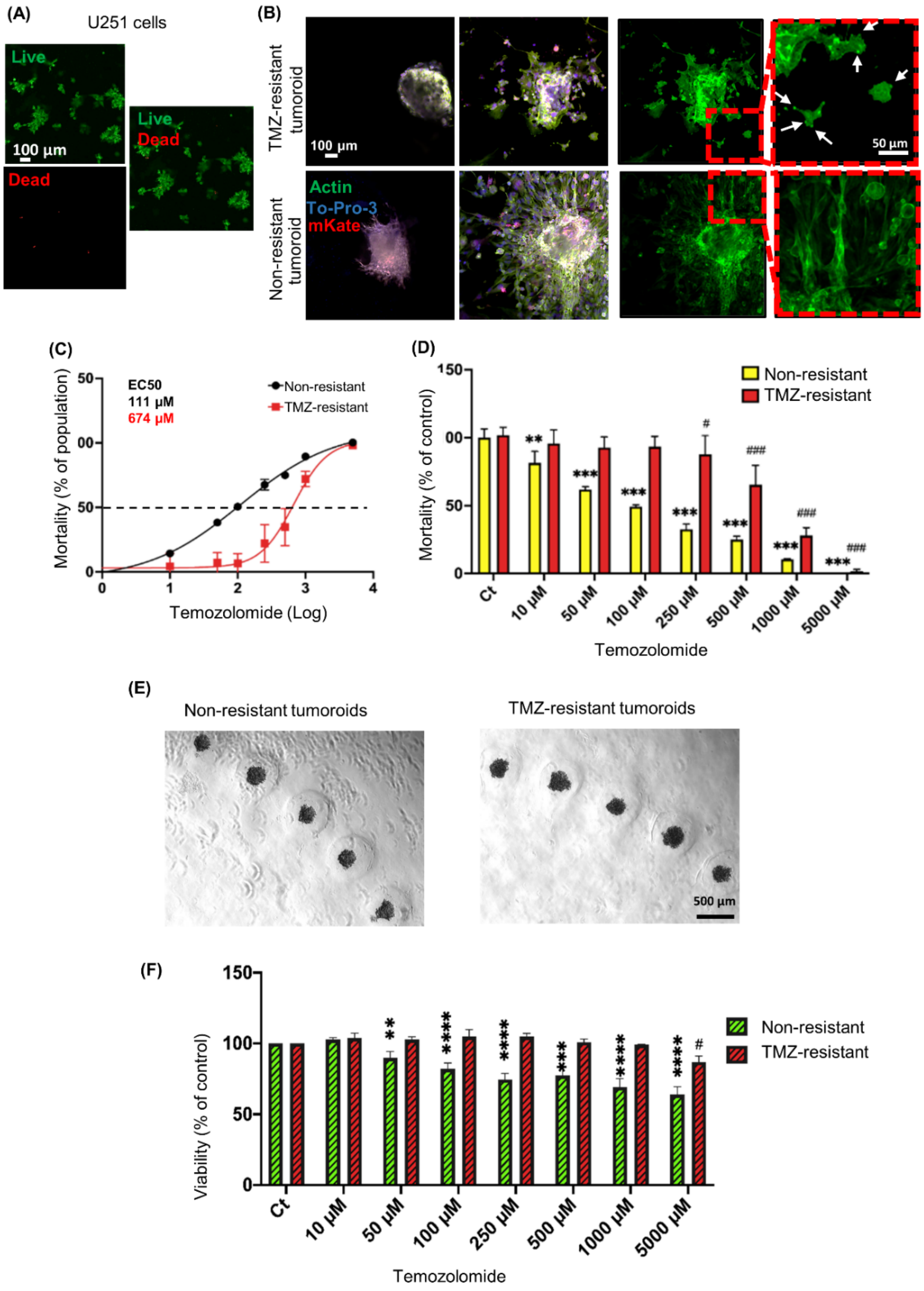


**Figure S1**. (A) Live/dead assay of U251 cells after 4 days encapsulation in CH. (B) Confocal imaging of actin filaments, visualized with phalloidin staining, comparing the invasion pattern of non-resistant *vs.* TMZ-resistant tumoroids. (C) EC50 and fold resistance of non-resistant and TMZ-resistant cells. (D) Viability assay of non-resistant and TMZ-resistant cells treated with 0-5000 µM of TMZ for 96h. (E) Formation of tumoroids. (F) Viability assay of 3D non-resistant and TMZ-resistant tumoroids treated with 0-5000 µM of TMZ for 96h. N=3 biological independent experiment in (A). The values are mean ± SEM of n =15 biologically independent samples examined over three independent experiments in (D, F).

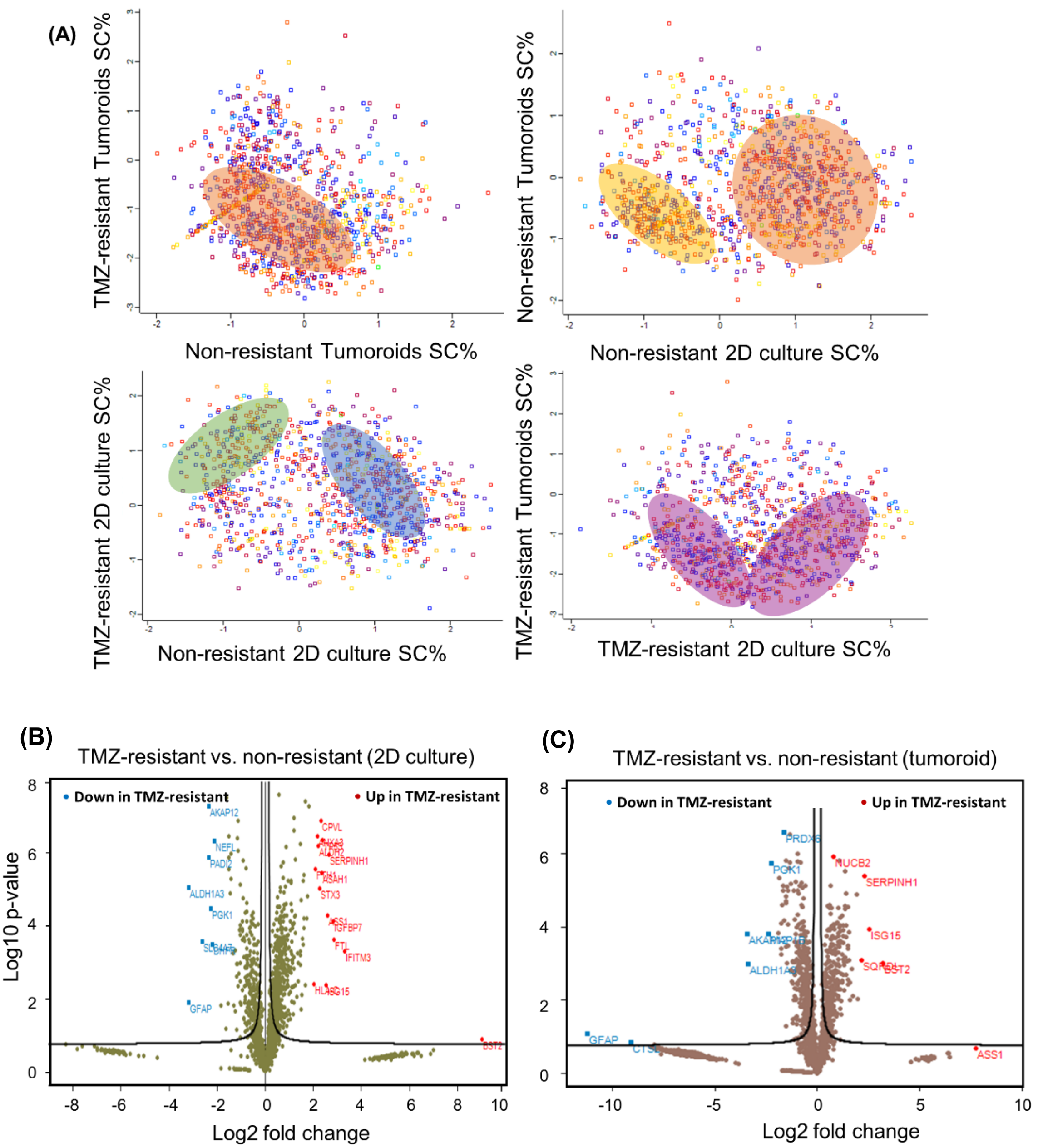


**Figure S2.** (A) Scatter plots for sequence converge percentage (SC%) in non-resistant and TMZ-resistant cells in 2D and 3D cultures. Volcano scatter plot of 2D (B) and 3D (C) non-resistant vs TMZ-resistant GB cells and tumoroids, respectively. The red or blue dots indicate proteins significantly ($q < 0.05$) upregulated (log2 fold change $\geq 0.5$) or downregulated (log2 fold change $\leq -0.5$) respectively. Proteins with more than 2-fold up or downregulation are annotated.

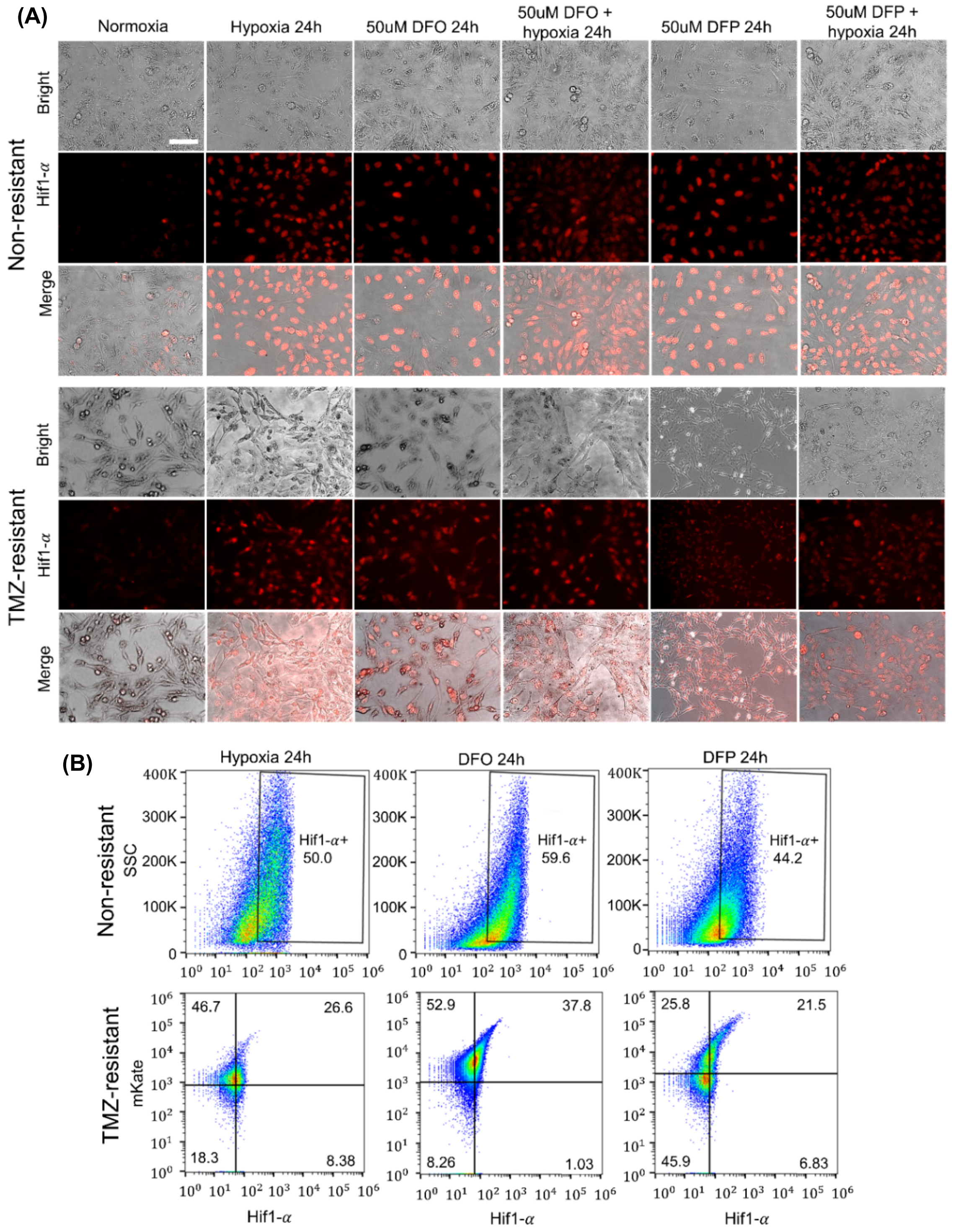


**Figure S3.** (A) Immunostaining of Hif1-α expression in U251 non-resistant and TMZ-resistant cells exposed to 24h of hypoxia, 50 μM DFO, 50 μM DFP, and combination of hypoxia and either DFO or DFP. (B) Semi-quantification of Hif1-α expression revealed that non-resistant cells exhibited higher accumulation of Hif1-α compared to TMZ-resistant cells in all treatment conditions. DFO treated non-resistant and TMZ-resistant cells expressed higher level of Hif1-α (%59, %37), respectively, compared to both DFP treatment (%40, %21) and hypoxia incubation (%50, 26%).

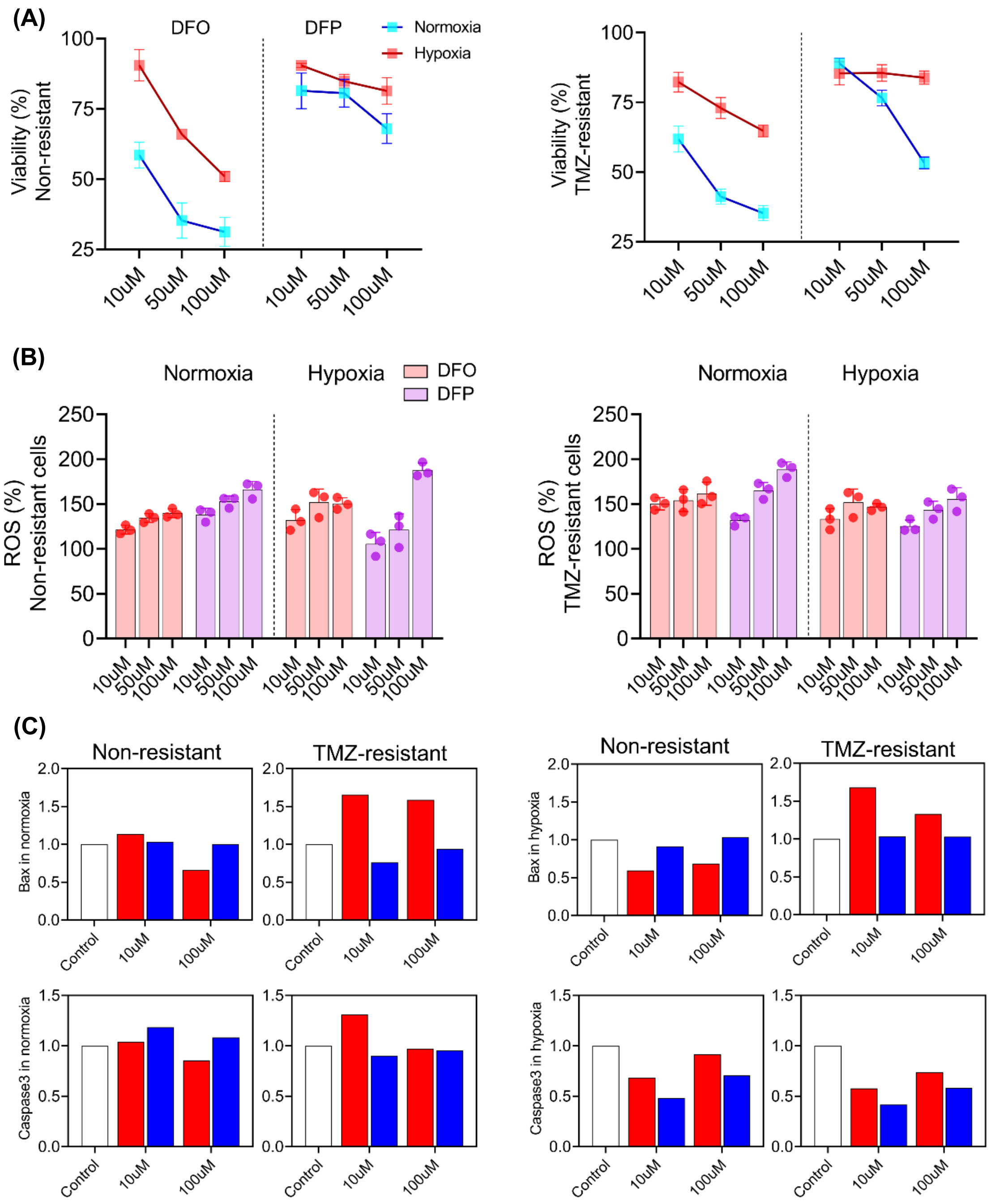


**Figure S4.** (A) Iron chelators reduced viability of non-resistant and TMZ-resistant cells after 72h, showing significant reduction in response to DFO but less sensitivity to DFP, with hypoxia counteracting the viability reduction. (B) Iron chelators induced ROS generation after 72h with hypoxia reducing the effect of DFP treatment in both non-resistant and TMZ-resistant cells. N=3 biological independent experiments and statistically significant at a P-value $< 0.05$. (C) Densitometry analysis of WB showed bax expression was increased only in response DFO in normoxic and hypoxic TMZ-resistant cells. However, bax was not increased in NR cells under any treatment. Although neither of the chelators showed significant increase in expression of caspase3, it was reduced in hypoxia.
